# Pitfalls in Post Hoc Analyses of Population Receptive Field Data

**DOI:** 10.1101/2020.12.15.422942

**Authors:** Susanne Stoll, Elisa Infanti, Benjamin de Haas, D. Samuel Schwarzkopf

## Abstract

Data binning involves grouping observations into bins and calculating bin-wise summary statistics. It can cope with overplotting and noise, making it a versatile tool for comparing many observations. However, data binning goes awry if the same observations are used for binning (selection) and contrasting (selective analysis). This creates circularity, biasing noise components and resulting in artifactual changes in the form of regression towards the mean. Importantly, these artifactual changes are a statistical necessity. Here, we use (null) simulations and empirical repeat data to expose this flaw in the scope of post hoc analyses of population receptive field data. In doing so, we reveal that the type of data analysis, data properties, and circular data cleaning are factors shaping the appearance of such artifactual changes. We furthermore highlight that circular data cleaning and circular sorting of change scores are selection practices that result in artifactual changes even without circular data binning. These pitfalls might have led to erroneous claims about changes in population receptive fields in previous work and can be mitigated by using independent data for selection purposes. Our evaluations highlight the urgency for us researchers to make the validation of analysis pipelines standard practice.

**Graphical abstract:** 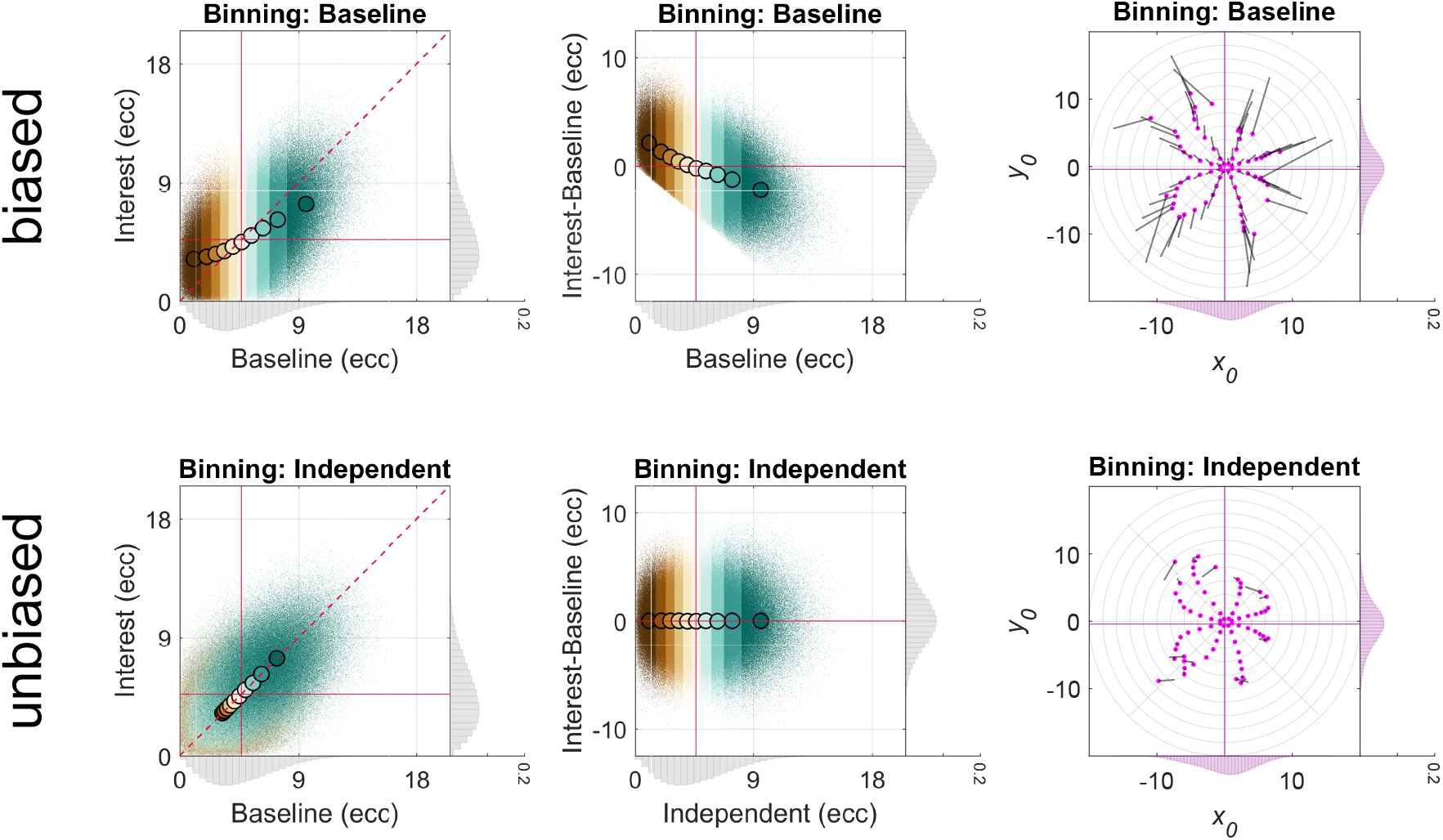

**Highlights:**

- Circular data binning produces artifactual changes in the form of regression towards the mean
- Analysis type, data properties, and circular data cleaning shape these artifactual changes
- Circular data cleaning and sorting produce artifactual changes even without circular data binning

## 1. Introduction

Data binning refers to grouping observations into bins or subgroups and calculating bin-wise summary statistics, such as the arithmetic mean. It is often applied to large datasets in order to prevent overplotting and control noise. As such, data binning has become commonplace in population receptive field (pRF) modeling (Dumoulin and Knapen, 2018; Dumoulin and Wandell, 2008), where researchers are commonly interested in comparing visual field maps with thousands of observations between different (experimental) conditions. However, pRF modeling is only one out of several research areas where some form of differential data binning has been adopted (e.g., Gignac and Zajenkowski, 2020; Holmes, 2009; Kriegeskorte et al., 2009; Preacher et al., 2005; Shanks, 2017).

Although data binning can help us see an overall pattern in the face of an abundance of details, it goes awry if the same observations are used for *binning* (selection) and *contrasting* (selective analysis). This is because dipping into noise-tainted data (i.e., most data) more than once violates assumptions of independence, favoring some noise components over others and eventually biasing descriptive and inferential statistics (Kriegeskorte et al., 2009). As such, double-dipping in data binning prevents us from – amongst other things – controlling for *regression towards the mean* (e.g., Galton, 1886; Gignac and Zajenkowski, 2020; Holmes, 2009; Makin and De Xivry, 2019; Shanks, 2017; Stigler, 1997).

Regression towards the mean is a statistical artifact occurring when two variables are imperfectly correlated (e.g., due to random noise^1^). In this case, extreme observations for one variable will on average be less extreme for the other^2^ (e.g., Campbell and Kenny, 1999; Cohen et al., 2003; Galton, 1886; Shanks, 2017; Stigler, 1997). The magnitude of regression towards the mean tends to be higher the lower the correlation between the variables (e.g., Campbell and Kenny, 1999, for systematic simulations, see Holmes 2009).

Double-dipping and/or regression towards the mean are of particular concern in what is known as *post hoc subgrouping* (Preacher et al., 2005), *post hoc data selection* (Shanks, 2017), and *extreme groups approach* (Preacher et al., 2005), all of which can be considered as subtypes of data binning. Post hoc subgrouping refers to collecting two measures, defining extreme subgroups post hoc using one measure (e.g., the lower and upper quantile), and then performing statistics on these measures for the extreme subgroups (Preacher et al., 2005). Post hoc data selection is similar but involves only one extreme subgroup (Shanks, 2017). Both of these practices are different from the extreme groups approach, where extreme subgroups are selected a priori based on one measure; that is, without collecting the whole range of the other measure (Preacher et al., 2005). Here, we focus on a post hoc scenario where essentially all subgroups are considered, not just the extreme ones (see also Gignac and Zajenkowski, 2020; Holmes, 2009). We label this procedure including its subtypes *post hoc binning analysis*.

An intuitive way to think about the link between double-dipping, regression towards the mean, and post hoc binning are repeat data. Assume we measure body weight in a population of adults twice – Today and Tomorrow (see endpoints of colorful lines, Figure 1; 1^st^ column). Further assume that any weight we measure involves a *permanent* and a *transient* component (true value + random noise). When determining Today’s and Tomorrow’s overall mean weight, all things being equal, the permanent component persists and the transient component cancels out (see red horizontal lines, Figure 1, 1^st^ column). However, this is not the case when we select adults with extremely high measurements for Today (relative to the overall mean) and compare these measurements to Tomorrow’s in the same adults by calculating the means (see lines and dots in dark green color, Figure 1, 1^st^ column). This is because we used Today’s measurements twice: for selection (binning) and selective analysis (comparing bin-wise means). We therefore favored Today’s noise components over Tomorrow’s. Why is this? The noise components of our selection criterion are not independent of the noise components of Today’s measurements. This renders the subgroup we selected Today on average heavier than it really is. This is not the case for Tomorrow’s measurements. As a result, Tomorrow’s measurements for this subgroup regress on average to Tomorrow’s overall mean (see dots in dark green color, Figure 1, 1^st^ column; for a similar example see Stigler, 1997). This artifactual change in average weight might look like a real phenomenon, although – of course – it is not.

The analysis we just performed can be regarded as an instantiation of post hoc data selection involving one extreme subgroup. If we additionally select a subgroup of adults with extremely low measurements for Today (see lines and dots in dark brown color, Figure 1, 1^st^ column), regression towards the overall mean from below occurs for this subgroup. Such an approach would qualify as post hoc subgrouping involving two extreme subgroups. If we incorporate additional less extreme subgroups, we perform a full-blown post hoc binning analysis (see lines and dots in various colors, Figure 1, 1^st^ column), where the bin-wise means for Tomorrow’s measurements regress towards the overall mean to various degrees. Importantly, this regression artifact is a statistical necessity not hinging upon body weight data. Once we use Independent data for binning purposes (e.g., body weight measurements collected for the day after tomorrow), we break the circularity, and the regression artifact disappears (Figure 1, 2^nd^ column).

**Figure 1.**
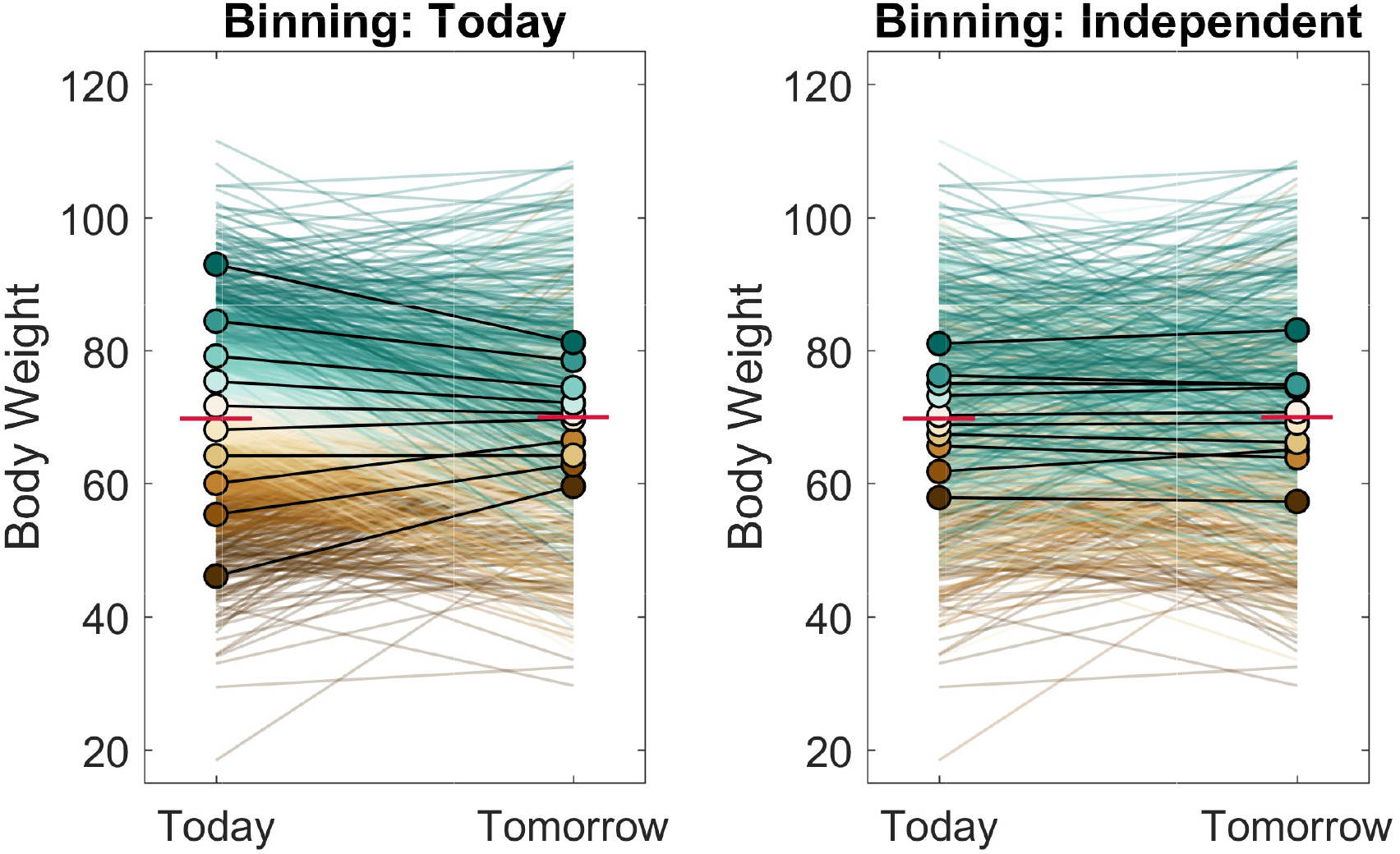
Simulated post hoc binning analysis on fictive body weight data. Bin-wise fictive body weight data and means for Today and Tomorrow in the same group of adults and different data binning scenarios. Body weight data for Today and Tomorrow were either binned according to body weight data for Today (1^st^ column) or an Independent test occasion (2^nd^ column). Fictive body weight data were simulated by sampling the body weight of 1000 adults from a Gaussian distribution (*M* = 70 kg; *SD* = 10 kg) and disturbing each adult’s body weight with random Gaussian noise (*SD* = 10 kg), separately for each test occasion (Today, Tomorrow, and Independent). The red horizontal lines indicate the location of the overall mean for Today and Tomorrow. Dark brown colors correspond to lower and dark blue-green colors to higher decile bins. The endpoints of the colorful lines represent individual data points and the colorful dots with the black outline bin-wise means. Note that the graphs displayed here are referred to as Galton squeeze diagrams (Campbell and Kenny, 1999; Galton, 1886; Shanks, 2017).

How does all of this relate to post hoc analyses involving pRF data? Imagine we conduct a retinotopic mapping experiment (Dumoulin and Wandell, 2008), where we estimate pRF position and pRF size for each voxel in the visual brain under a *Baseline* condition as well as a condition of *Interest* (see Figure 2 for a single pRF). We can think of the Interest and Baseline conditions as repeat data (e.g., Benson et al., 2018; Senden et al., 2014; van Dijk et al., 2016), different attention conditions (e.g, de Haas et al., 2014, 2020; Klein et al., 2014; van Es et al., 2018; Vo et al., 2017), mapping sequences (e.g., Binda et al., 2013; Infanti and Schwarzkopf, 2020), mapping stimuli (e.g., Alvarez et al., 2015; Binda et al., 2013; Le et al., 2017; Yildirim et al., 2018), magnetic field strengths (e.g., Morgan and Schwarzkopf, 2020), scotoma conditions (e.g., Barton and Brewer, 2015; Binda et al., 2013; Haak et al., 2012; Prabhakaran et al., 2020), and pRF modeling techniques (e.g., Carvalho et al., 2020) – to name but a few examples. Similarly, apart from visual scenarios, we can also interpret the Baseline and Interest condition as adaptation conditions (e.g., Tsouli et al., 2021), different finger movements (e.g., Schellekens et al., 2018), or uni- and multisensory conditions (see Holmes, 2009, for a discussion on non-pRF work).

**Figure 2.**
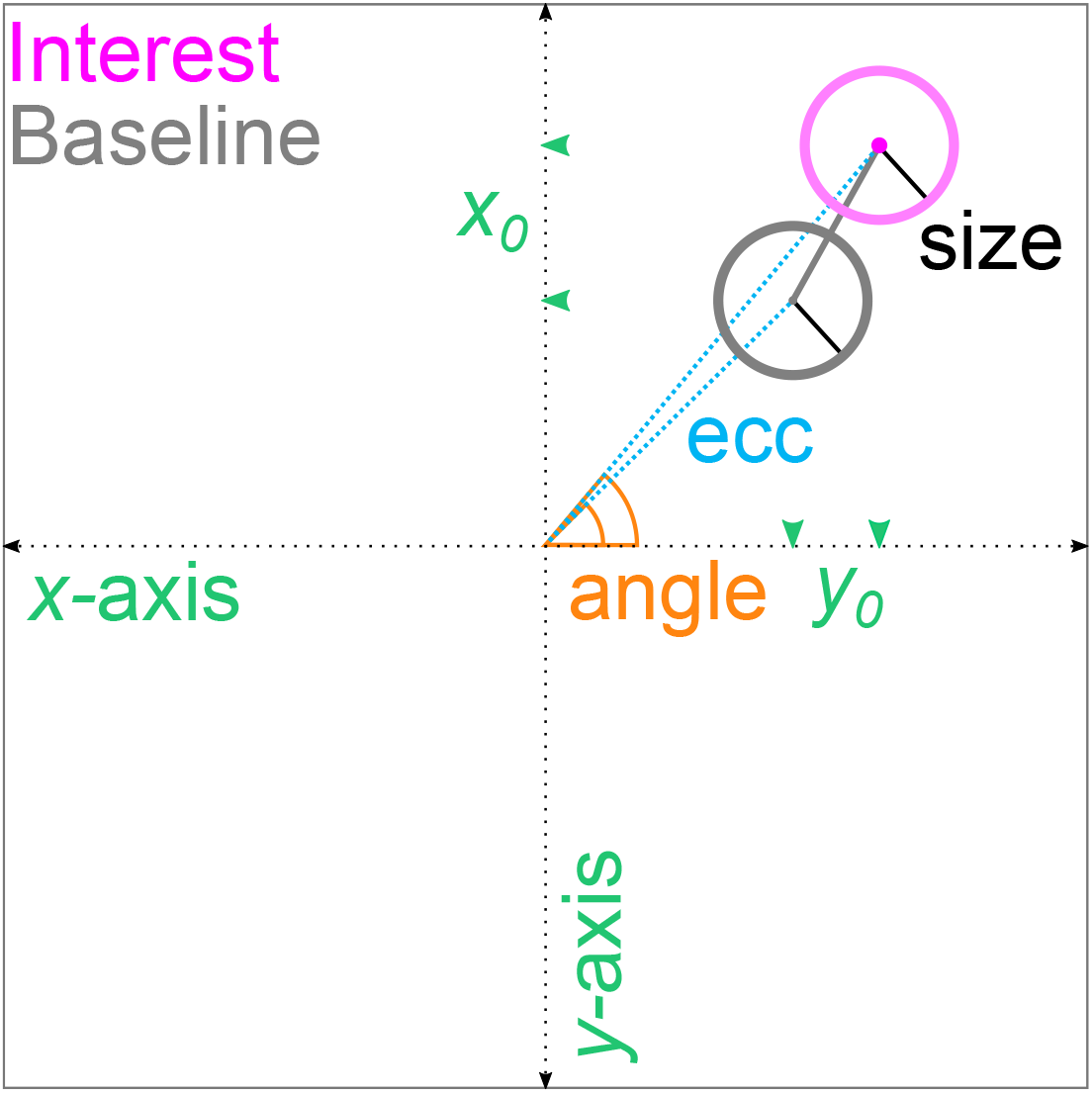
Population receptive field estimates. The large black square outline represents a cutout of the visual field and the black dashed arrows a Cartesian coordinate system. The two circles represent a pRF that changes its position (gray solid line) in an Interest (magenta) compared to a Baseline (gray) condition. The pRF was modeled as a 2D Gaussian function. The center of the 2D Gaussian (midpoint of the gray and magenta circles) represents the position of the pRF. PRF position can be expressed in terms of *x*_0_ and *y*_0_ coordinates (green arrow heads) or eccentricity (blue dashed line) and polar angles (orange solid line). Eccentricity corresponds to the Euclidean distance between the center of gaze (origin) and the center of the 2D Gaussian. Polar angle corresponds to the counter-clockwise angle running from the positive *x*-axis to the eccentricity vector. The standard deviation of the Gaussian (1*σ*; black solid line) represents pRF size. Both pRF position and size are typically expressed in degrees of visual angle. Polar angles are typically expressed in degrees. Ecc = Eccentricity. pRF = Population receptive field.

As a pRF model, we adopt a 2D Gaussian, where pRF position represents the center of a pRF in visual space (the center of the Gaussian) and pRF size its spatial extent (the standard deviation of the Gaussian; see Figure 2). We then fit this model to the voxel-wise brain responses we measured in the retinotopic mapping experiment (Dumoulin and Wandell, 2008). To compare pRF positions in the Interest and Baseline condition voxel-by-voxel, we bin the pRF positions from both conditions according to the pRF positions from the Baseline condition. Subsequently, we quantify for each voxel the position shift from the Baseline to the Interest condition (see Figure 2 for a single pRF). Finally, we calculate the bin-wise mean shift. This is equivalent to calculating the bin-wise simple means for each condition and comparing them subsequently.

Either way, by adopting such a post hoc binning analysis, we essentially assume that binning voxels according to pRF positions from the Baseline condition and aggregating them subsequently for this condition ensures that bin-wise noise components are unbiased on average (see also Shanks, 2017). This, however, is not the case. The underlying reason is the same as for our body weight analysis further above: we dipped into the Baseline condition twice, namely to define bins (selection) and to estimate bin-wise means for further comparison (selective analysis). This circularity leads to a favoring of noise components, skewing the bin-wise means in the Baseline condition and eventually resulting in regression towards the overall mean for the bin-wise means of the Interest condition.

Here, we expose and explore this flaw in the scope of post hoc analyses of pRF data using (null) simulations and empirical repeat data from the Human Connectome Project (HCP; Benson et al., 2018, 2020). Unlike empirical data, simulations allowed us to separate true values from noise components. They also provided an excellent test bed for determining that the type of data analysis (change scores or simple scores, 1D or 2D binning, equidistant or decile binning), data properties (presence or absence of heteroskedasticity or a true effect) and additional circular selection practices (presence or absence of circular data cleaning) influence the appearance of the regression artifact. Moreover, they allowed us to pinpoint that circular data cleaning and circular sorting of change scores represent selection practices that yield artifactual changes even without circular data binning. Unlike empirical data from different experimental conditions, repeat data permitted us to assume a null effect between conditions, allowing for more straightforward conclusions about any systematic differences we might observe.

## 2. Methods

### 2.1. Post hoc binning using simulated data

For the post hoc binning analysis involving simulations, we used an empirical V1 visual field map of a single human participant as a basic data distribution. This map originated from a functional magnetic resonance imaging experiment (fMRI) aimed at mapping pRFs under different attention conditions using a drifting bar stimulus (2 sessions each with 4 runs per condition). One of these conditions was selected for simulation purposes. The maximal eccentricity of the mapping area subtended 8.5 degrees of visual angle (dva). We fit a 2D Gaussian function to preprocessed fMRI responses projected onto the cortical surface. For each vertex (gray matter node on the cortical surface), we obtained 6 estimates: pRF position (*x*_0_ and *y*_0_ coordinates), pRF size (*σ*), pRF baseline (*β*_0_), pRF amplitude (*β*_1_), and goodness-of-fit (*R*^2^). We first smoothed the resulting parameter maps and delineated V1 hemifield maps manually (for more details, see Supplementary methods, 1. Retinotopic mapping experiment). We then pooled the *x*_0_ and *y*_0_ coordinates across V1 hemifield maps and removed empty data points.

#### 2.1.1. 1D post hoc binning analysis on eccentricity

To uncover the regression artifact, we first simulated a simplified contrast scenario with a null effect. To this end, we disturbed the *x*_0_ and *y*_0_ coordinates (Figure 2) 200 times with random Gaussian noise (*SD* = 2 dva). We repeated this to generate a *Baseline*, *Interest*, and *Independent* condition. We then converted the *x*_0_ and *y*_0_ coordinates to eccentricity values (Figure 2), as is often done in the pRF literature (see Figure S1 for interpretational difficulties with eccentricity when it comes to position shifts). This resulted in a gamma-like eccentricity distribution. Lastly, we binned the eccentricity values in the Baseline and Interest condition according to the eccentricity values of any of the 3 conditions using deciles and calculated the bin-wise means^3^. A schematic workflow of this simulated 1D post hoc binning analysis can be found in Figure 3. Bin-wise eccentricity means were visualized as a color-coded scatter plot along with individual observations per bin and marginal histograms (bin width = 0.5 dva) reflecting the simulated distributions.

**Figure 3.**
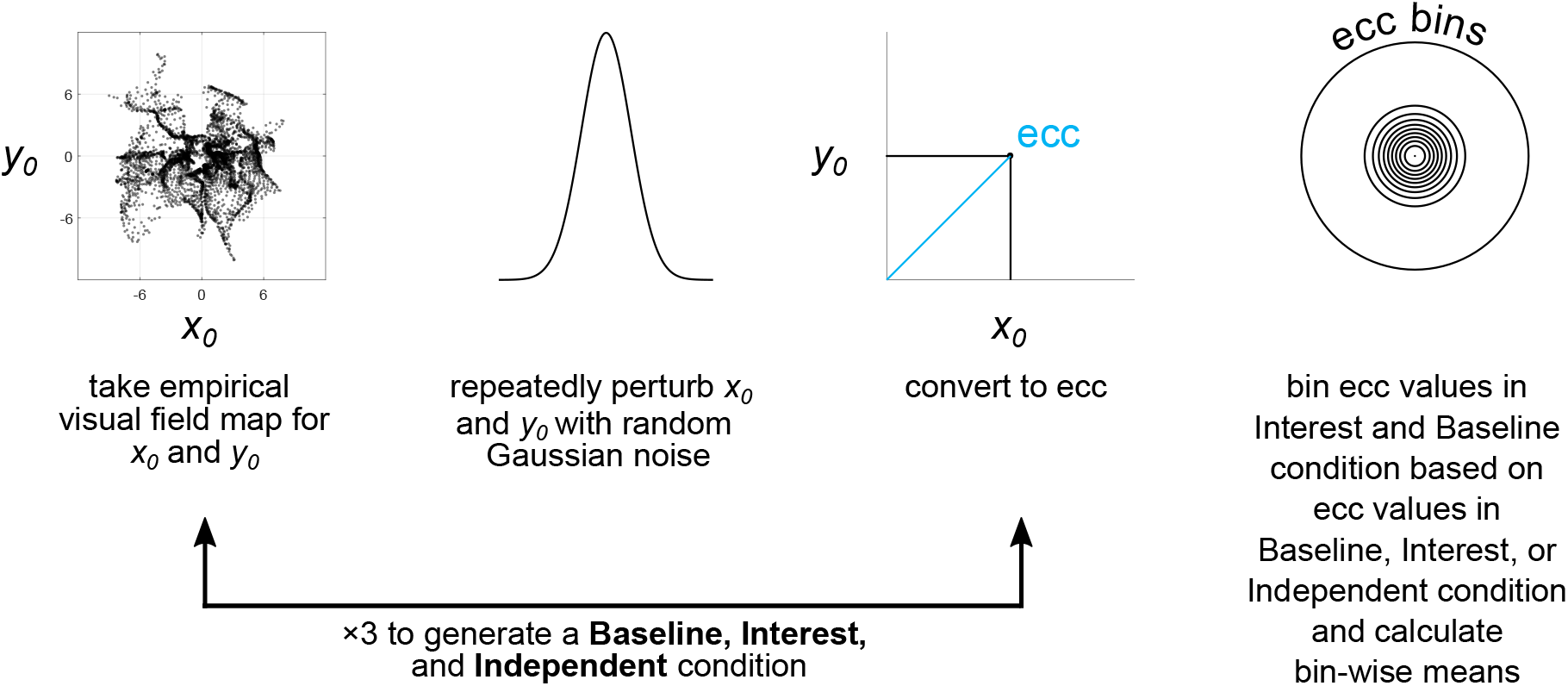
Schematic workflow of 1D post hoc binning analysis on simulated eccentricity data | Null effect. Ecc = Eccentricity.

Building upon the simulated null effect, we performed the 1D post hoc binning analysis on 4 more simulation cases: a null effect with condition cross-thresholding based on the Baseline condition, a null effect with condition cross-thresholding based on both the Baseline and Interest condition, a null effect with eccentricity-dependent noise, and a true effect. We use the term ‘condition cross-thresholding’ to refer to the pair-wise or list-wise deletion of data points across experimental conditions (see below). The selected simulation cases reflect analysis practices and data properties we consider characteristic of pRF studies. For all simulation cases, the Independent condition consisted of a second draw (resample) of the Baseline condition. Moreover, to ensure reproducibility and comparability, all simulation cases were based on the same seed for random number generation. However, our conclusions do not depend on the choice of seed for random number generation.

For the simulation cases involving condition cross-thresholding, we removed simulated observations falling outside a certain eccentricity range (*≥* 0 and *≤* 6 dva) in the Baseline or Baseline and Interest condition from all conditions (i.e., Baseline, Interest, and Independent). For the simulation case involving eccentricity-dependent noise, we used a small standard deviation (*SD* = 0.25 dva) of random Gaussian noise to disturb empirical observations with smaller eccentricities (*≥* 0 and *<* 3 dva) and a larger standard deviation (*SD* = 2 dva) to disturb empirical observations with larger eccentricities (*≥* 3 dva). For the simulation case involving a true effect, we induced a radial increase in eccentricity of 2 dva in the Interest condition.

Apart from simple bin-wise means, we performed the 1D post hoc binning analysis also on change scores. The change scores were obtained by subtracting individual simulated observations or means in the Baseline condition from those in the Interest condition. Both simple means and mean change scores have been used for post hoc binning in previous pRF studies (e.g., Barton and Brewer, 2015; Binda et al., 2013; Carvalho et al., 2020; Haak et al., 2012; Yildirim et al., 2018; de Haas et al., 2014, 2020; Prabhakaran et al., 2020; Tsouli et al., 2021). Similarly, we repeated the binning analysis using equidistant instead of decile binning. To this end, we used a constant bin width of 1.75 dva and an overall binning range of 0 to 19.25 dva eccentricity. Unlike equidistant binning, decile binning ensures a roughly equal number of data points in each bin, which facilitates the interpretation of results. However, we consider equidistant binning as the most common binning type in the pRF literature. For both the change score analysis and equidistant binning, we used the simulation case involving a null effect as a data basis.

#### 2.1.2. 2D post hoc binning analysis on x_0_ and y_0_

Apart from the 1D binning analysis on eccentricity, we also conducted a 2D binning analysis on the simulated *x*_0_ and *y*_0_ values. To this end, we converted the *x*_0_ and *y*_0_ values to polar coordinates, that is, polar angle and eccentricity (Figure 2). We then binned the *x*_0_ and *y*_0_ values in the Baseline or Interest condition according to their polar coordinates in the Baseline, Interest, or Independent condition using equidistant bins and calculated the bin-wise *x*_0_ and *y*_0_ means for each condition. The condition-wise means were visualized as vector graphs along with marginal histograms (bin width = 0.5 dva) illustrating the simulated distributions. Vector graphs have been used in prior pRF work (e.g., Klein et al., 2014; van Es et al., 2018; Vo et al., 2017). The 2D binning analysis was performed for all aforementioned simulation cases. The polar angle bins ranged from 0° to 360° with a constant bin width of 45°. The eccentricity bins ranged from 0 to 22 dva (for the simulation case involving a true effect) or from 0 to 20 dva (for all other simulation cases) with a constant bin width of 2 dva.

### 2.2. Post hoc binning using empirical repeat data

For the post hoc binning analysis on repeat data, we used publicly available pRF estimates from the HCP 7 T Retinotopy Dataset (Benson et al., 2018, 2020). These estimates stem from a split-half analysis where a 2D isotropic Gaussian with a subadditive exponent (Kay et al., 2013) was fit to fMRI time series from the first and second half of 6 pRF mapping runs. For each half, 6 estimates were obtained for each grayordinate (vertex; https://wiki.humanconnectome.org/display/WBPublic/Workbench+Glossary), that is, pRF polar angle, pRF eccentricity, pRF size, pRF gain, percentage of *R*^2^, and mean signal intensity. The maximal eccentricity of the mapping area subtended 8 dva. For further details, see Benson et al. (2018).

Following Benson et al. (2018), we analyzed complexes of visual areas across hemispheres for the 25^th^ and 75^th^ percentile participants of the *R*^2^ distribution using delineations from Wang et al.’s (2015) atlas. Benson et al. (2018) generated the *R*^2^ distribution by calculating the median *R*^2^ for each participant across grayordinates from both cortical hemispheres within all areas of Wang et al.’s (2015) atlas. For our purposes, we focused on the posterior complex (V1-V3) and the dorsal complex (V3A/B and IPS0–5), as those came with a larger number of available data points (which was, amongst other things, necessary to perform the 2D post hoc binning analysis and generate vector graphs).

To obtain *x*_0_ and *y*_0_ values, polar angle and eccentricity estimates were converted to Cartesian coordinates. The eccentricity, *x*_0_, and *y*_0_ values of the first half were used as a Baseline condition and those of the second half as an Interest condition. Similar to the simulation-based analyses, binning was either based on the Interest or Baseline condition and bin-wise means were calculated. Moreover, binning was either performed without or with condition cross-thresholding. As for the latter case, we removed observations outside a certain eccentricity range (*≥* 0 and *≤* 8 dva) or below a certain *R*^2^ cut-off (*≤* 2.2%) in the Baseline or Baseline and Interest condition from both conditions. The *R*^2^ cut-off was adopted from Benson et al. (2018).

We then performed a 1D binning analysis on eccentricity and a 2D binning analysis on *x*_0_ and *y*_0_ as we did for the simulated data. However, here, the eccentricity bins for the 2D analysis ranged from 0 to 18 dva with a constant bin width of 2 dva. All binning analyses and visualizations (including those on simulated data) were implemented in Matlab 2016b (9.1; https://uk.mathworks.com/) using custom code (Data and code availability). The color scheme used for color-coding was an adapted version of the BrBG palette from ColorBrewer (2.0; Brewer et al., 2021) retrieved via R (3.5.3; R Core Team, 2018) and the package RColorBrewer (1.1-2; Neuwirth, 2014).

## 3. Results and discussion

### 3.1. The many faces of regression towards the mean and other problems

To expose the regression artifact, we repeatedly perturbed the *x*_0_ and *y*_0_ values of an empirical visual field map with random Gaussian noise to generate a Baseline and Interest condition. We then converted the *x*_0_ and *y*_0_ values to eccentricity. Subsequently, we binned the eccentricity values of either condition according to eccentricity values in the Baseline condition using deciles and calculated bin-wise means. The bin-wise means from both conditions were plotted against one another along with individual observations per bin and marginal histograms reflecting the simulated distributions^4^ (Figure 4, 1^st^ column). Since there was no true difference between conditions, the bin-wise means should lie on the identity line. Contrary to this prediction, they systematically diverged from the identity line. Strikingly, when using the Interest instead of the Baseline condition for binning, this systematic pattern of divergence flipped (Figure 4, 2^nd^ column). This bidirectionality is a typical sign of regression towards the mean (Campbell and Kenny, 1999; Shanks, 2017) and due to circularity. This leads to asymmetric bins (see bin-wise ranges of observations for the Baseline and Interest condition, Figure 4, 1^st^ and 2^nd^ columns) and on average biases bin-wise noise components for the condition that was used for contrasting and binning (henceforth *circular* condition). On the contrary, for the other condition (henceforth *non-circular* condition), this is not the case.

**Figure 4.**
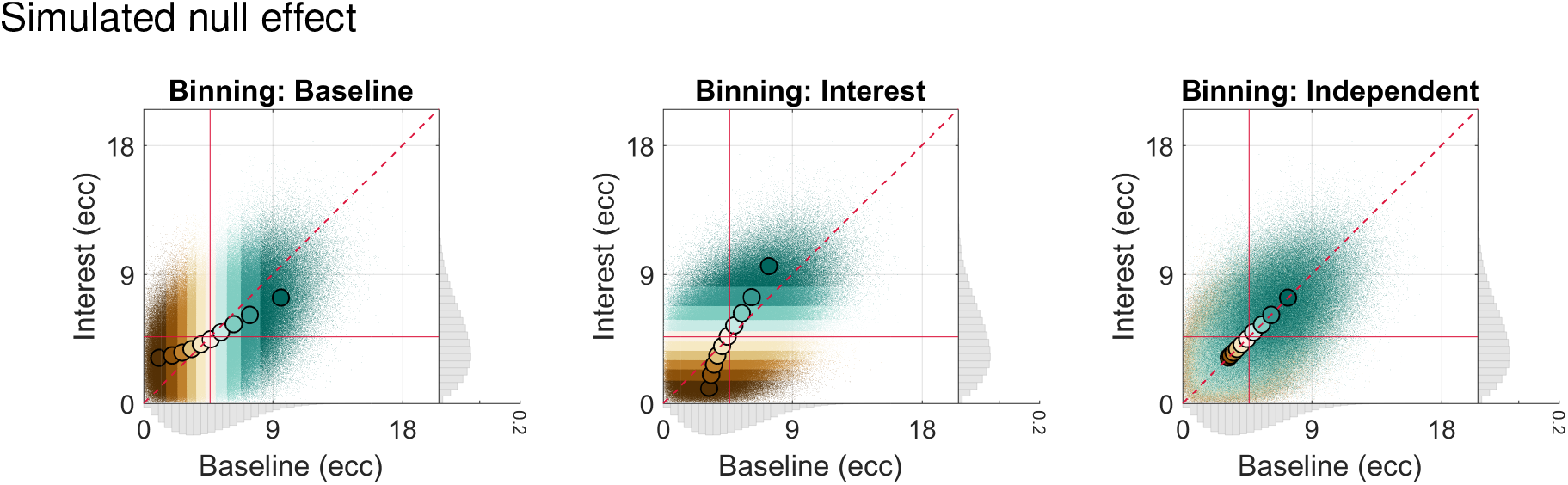
Simulated 1D post hoc binning analysis on eccentricity | Null effect. Bin-wise eccentricity values and means in the Interest and Baseline condition for a simulated null effect and different data binning scenarios. The eccentricity values in the Baseline and Interest condition were either binned according to eccentricity values in the Baseline (1^st^ column), Interest (2^nd^ column), or an Independent condition (equivalent to repeat data of the Baseline condition; 3^rd^ column). The gray marginal histograms (bin width = 0.5 dva; *y*-axis: relative frequency) show the simulated eccentricity distributions for each condition, obtained by repeatedly disturbing the *x*_0_ and *y*_0_ values of an empirical visual field map with random Gaussian noise (*SD* = 2 dva) and subsequently converting them to eccentricity values. Note that the range of the marginal *y*-axis is the same for all histograms. The red crosshair indicates the location of the overall mean for the Interest and Baseline condition. The red dashed line corresponds to the identity line. Dark brown colors correspond to lower and dark blue-green colors to higher decile bins. The smaller colorful dots represent individual data points and the larger colorful dots with the black outline bin-wise means. The maximal eccentricity of the stimulated visual field area subtended 8.5 dva. Dva = Degrees of visual angle. Ecc = Eccentricity

The skew in average noise renders the bin-wise eccentricity means of the circular condition more extreme, especially for lower and higher decile bins. As a result, the bin-wise eccentricity means for the non-circular condition regress – by statistical necessity – to the overall mean^5^ for this condition (red crosshair); that is, they are less extreme. This becomes clear when looking at the different ranges of bin-wise means for the circular and non-circular conditions (Figure 4, 1^st^ and 2^nd^ columns). If the Interest condition is then contrasted to the Baseline condition, a mean increase in eccentricity for lower deciles and a mean decrease for higher deciles or vice versa occurs, depending on whether the data are binned on the Baseline or Interest condition (Figure 4, 1^st^ and 2^nd^ columns). This artifact arises because we did not always use independent conditions for binning and contrasting; that is, conditions with independent noise components.

Apart from simple means (e.g., Binda et al., 2013; Carvalho et al., 2020; Haak et al., 2012; Yildirim et al., 2018), post hoc binning analyses have also been performed on change scores in previous pRF studies (e.g., Barton and Brewer, 2015; de Haas et al., 2014, 2020; Prabhakaran et al., 2020; Tsouli et al., 2021). Here, the difference between the Interest and Baseline condition is typically plotted against the binning (i.e., circular) condition (Figure 5, A., 1^st^ and 2^nd^ columns). Consequently, the bin-wise means now regress to the overall mean of the change score distribution (see also Gignac and Zajenkowski, 2020; Holmes, 2009) and bin-wise noise components are neither unbiased for the change scores nor the binning conditions. This is because the noise components of the change scores are not independent of those in either binning condition. What is more, scatter plots of change scores disguise important aspects readily available with scatter plots of simple scores. Specifically, they prevent us from directly appreciating the larger bin-wise range of eccentricity means for the circular as compared to the non-circular condition (see explanations further above and compare Figure 5, A., and Figure 4, 1^st^ and 2^nd^ columns). This makes it difficult to spot the source of the problem graphically when only looking at a single plot. On the other hand, since both the *x*- and *y*-axis feature the Baseline or Interest condition and either of these conditions are used for data binning, the act of double-dipping becomes much more obvious.

**Figure 5.**
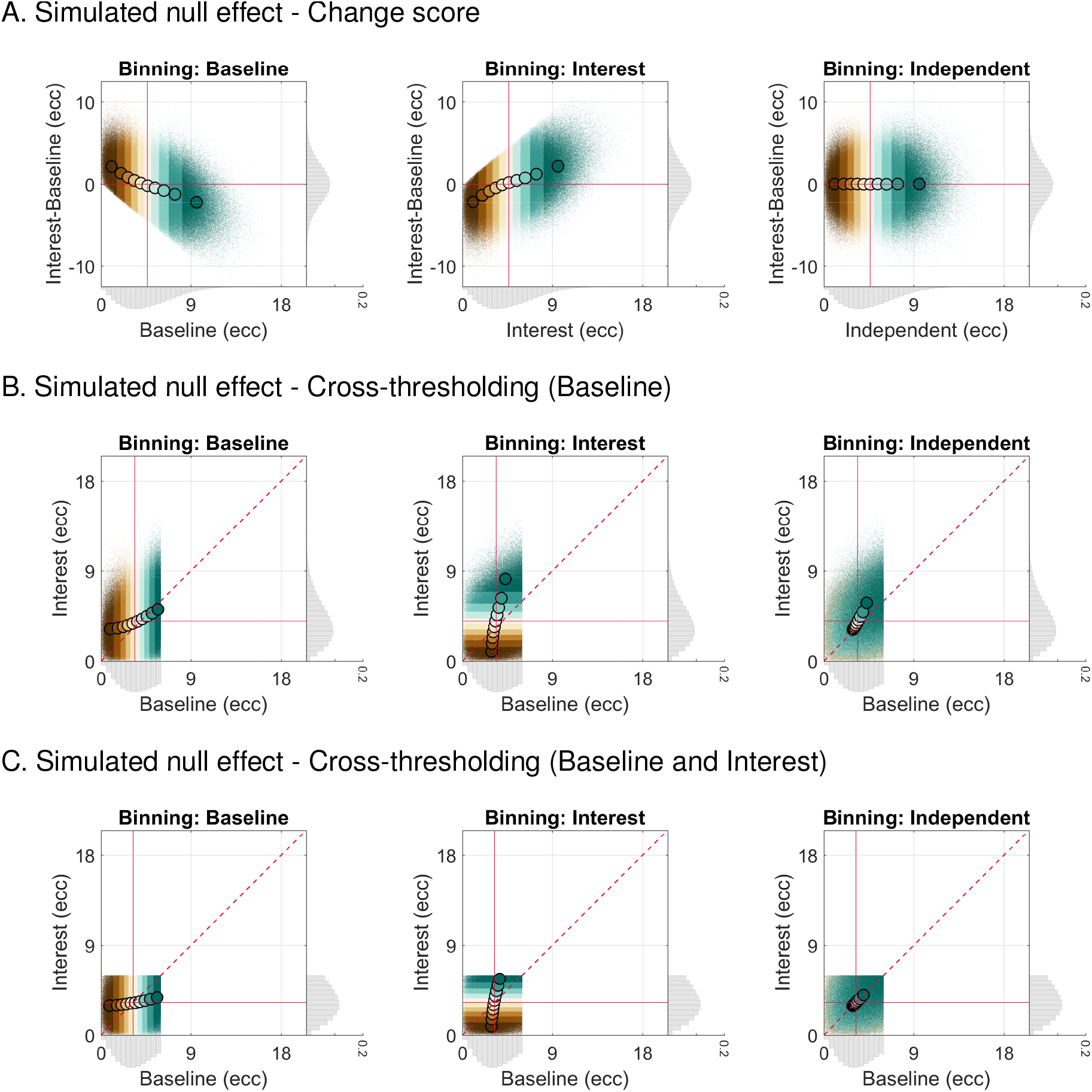
Simulated 1D post hoc binning analysis on eccentricity | Null effect — Change score and cross-thresholding. **A.** The same as in Figure 4, although here, the change score (Interest vs Baseline) is plotted against the respective binning condition. **B.** The same as in Figure 4, although here, condition cross-thresholding was applied, i.e., simulated observations falling outside a certain eccentricity range (*≥* 0 and *≤* 6 dva) in the Baseline condition were removed from all conditions. **C.** The same as in B., although here, condition cross-thresholding was based on both the Baseline and Interest condition. (Condition) cross-thresholding = The pair-wise or list-wise deletion of observations across conditions.

Critically, scattering change scores against one of the conditions involved in change score calculation also results in a biased visualization of individual change scores. This is because the noise components of the variables on the *x*- and *y*-axis are not independent, rendering this sorting procedure circular. When plotting individual change scores against the Baseline condition, this results in a downwards sloping data cloud, suggesting an effect although there is none (Figure 5, A., 1^st^ column). Why does this happen? Owing to noise, the change scores are more likely to be positive for lower Baseline eccentricities and negative for higher Baseline eccentricities (Figure 5, A., 1^st^ column). When plotting individual change scores against the Interest condition, the reverse is true (Figure 5, A., 2^nd^ column). This means visualizing or analyzing the data using such a circular sorting procedure is misleading irrespective of circular data binning (for more details on circular data sorting, see Holmes, 2009; Kriegeskorte et al., 2009).

The fact that circular sorting of change scores and circular data binning are separate issues can be further appreciated by imagining what happens when we plot the individual change scores against the Baseline condition, but bin on the Interest condition (instead of the Baseline condition as before). In this case, the individual change scores are sorted in a way (downwards sloping; just like in Figure 5, A., 1^st^ column) that is opposite to the trend implied by the bin-wise means (upwards sloping).

How the regression artifact induced by circular data binning manifests can change when data are thresholded across conditions, that is, deleted in a pair- or list-wise fashion (Figure 5, B. and C., 1^st^ and 2^nd^ columns). In fact, in the event of condition cross-thresholding, noise components are reshaped and might thus not necessarily be unbiased on average even for the non-circular condition (Figure 5, B., 2^nd^ column as well as Figure 5, C., 1^st^ and 2^nd^ columns). Condition cross-thresholding is common practice in the pRF literature where data are cleaned across conditions according to eccentricity, goodness-of-fit (*R*^2^), pRF size, missing data or other criteria from one or multiple conditions.

Here, we cross-thresholded the eccentricity values in the Interest and Baseline condition using the eccentricity values from the Baseline condition (Figure 5, B., 1^st^ and 2^nd^ columns) or both the Baseline and Interest condition (Figure 5, C., 1^st^ and 2^nd^ columns). This cross-thresholding procedure is circular whenever the noise components of the data used for cross-thresholding are not independent of the noise components of the data involved in contrasting. This is evidently true even without circular data binning. As such, the reason why the noise components in our cross-thresholding scenarios are sometimes biased even for the non-circular condition^6^ (Figure 5, B., 2^nd^ column as well as Figure 5, C., 1^st^ and 2^nd^ columns) is because we introduced another layer of circularity.

The fact that circular cross-thresholding and circular data binning are somewhat distinct but also highly similar issues can, for instance, be appreciated when comparing the overall instead of the bin-wise means. Without circular cross-thresholding, the overall mean in both the Baseline and Interest condition amounts to 4.66 dva (Figure 4, A., 1^st^ and 2^nd^ columns). With circular cross-thresholding based on the Baseline condition, the overall mean in the Baseline condition amounts to 3.40 dva, whereas it amounts to 3.97 dva in the Interest condition (Figure 5, B., 1^st^ and 2^nd^ columns). Here, the introduced bias for the Baseline condition can be appreciated by directly comparing the overall means in the Baseline and Interest condition. With circular cross-thresholding based on both the Baseline and Interest condition, the overall means in the Baseline and Interest condition amount to 3.24 dva and 3.25 dva, respectively (Figure 5, C., 1^st^ and 2^nd^ columns). Here, the introduced bias for the Baseline and Interest condition can be appreciated by comparing the overall means in these conditions to the overall mean of an Independent condition (retest of the Baseline condition) that was cross-thresholded based on both the Baseline and Interest condition. This overall mean amounts to 3.66. We will return to the usefulness of such an Independent condition further below (3.2. Potential mitigation strategies). In any case, circular cross-thresholding biases the overall means as compared to when no such thresholding is performed.

Importantly, however, only circular cross-thresholding based on the Baseline condition results in artifactual differences between the overall means. Why is this? Given that the level of noise in the Interest and Baseline condition was equivalent (2.1. Post hoc binning using simulated data), circular cross-thresholding based on both the Baseline and Interest condition on average skewed the noise components for these conditions similarly, resulting in biased overall means, but a valid difference of around 0 between them. However, as for empirical data, the assumption of equivalent noise levels can probably only be safely made for repeat data (and even then, this needs to be justifiable). In any case, conceptually, circular cross-thresholding without data binning can be regarded as a single bin or region-of-interest analysis (Kriegeskorte et al., 2009), essentially constituting another subtype of a post hoc binning analysis.

The appearance of the regression artifact arising from circular data binning can furthermore change when the level of noise depends on eccentricity – a property better known as *heteroskedasticity* (Figure 6, A., 1^st^ and 2^nd^ columns; see also Holmes, 2009). In fact, the case of eccentricity-dependent noise shows that the artifact can include some clear regression *away* from the mean – a phenomenon referred to as *egression*^7^ (Figure 6, A., 1^st^ and 2^nd^ columns; see e.g., Campbell and Kenny, 1999; Schwarz and Reike, 2018). Eccentricity-dependent noise might arise from fitting errors that differ across visual space. This could be due to partial stimulation of pRFs (especially near the edge of the stimulated mapping area), higher variability in pRF position estimates for wider pRFs as well as fluctuations in the signal-to-noise ratio of brain responses from the central to the peripheral visual field or as a result of manipulating attention.

The regression artifact due to circular data binning also manifested when simulating a true effect (Figure 6, B., 1^st^ and 2^nd^ columns). The same was true for equidistant binning (Figure 6, C., 1^st^ and 2^nd^ columns), which is frequently applied in the pRF literature. However, unlike decile binning (which we used further above), equidistant binning resulted in a lower number of observations for higher equidistant bins (due to the gamma-like eccentricity distribution; Figure 6, C., 1^st^ and 2^nd^ columns). Consequently, for higher equidistant bins, the skew in average noise for the circular condition was generally larger here (compare Figure 6, C., and Figure 4, 1^st^ and 2^nd^ columns). Similarly, for higher equidistant bins, noise components were not always unskewed on average for the non-circular condition (see Figure 6, C., 1^st^ and 2^nd^ columns, where the pattern of bin-wise means is not entirely mirror-symmetric). This is because for random noise to be unskewed on average, the number of observations needs to be sufficiently large.

**Figure 6.**
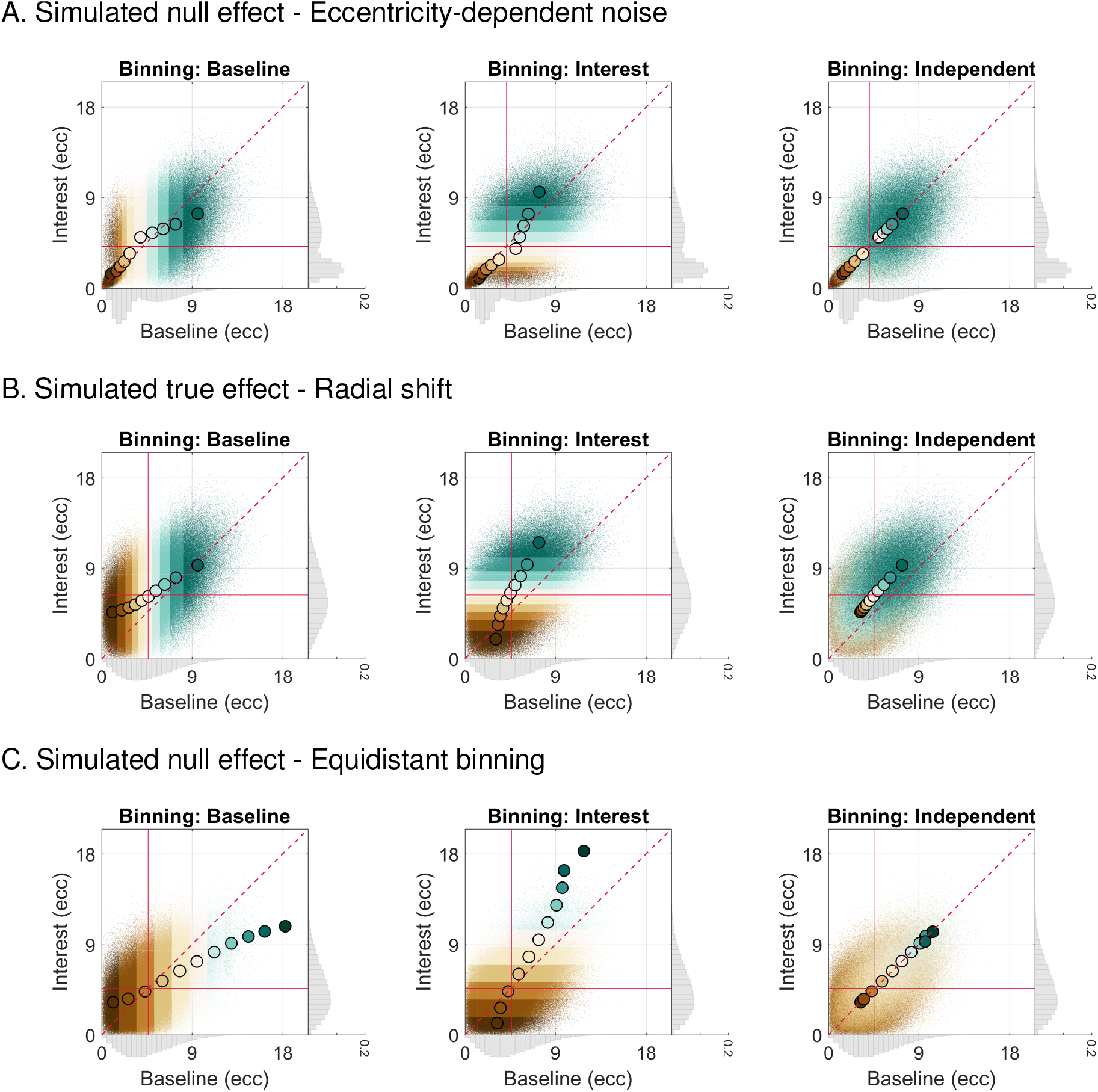
Simulated 1D post hoc binning analysis on eccentricity | Null or true effect — Eccentricity-dependent noise, radial shift, and equidistant binning. **A.** The same as in Figure 4, although here, original observations having smaller eccentricities (*≥* 0 and *<* 3 dva) were disturbed by random Gaussian noise with a smaller standard deviation (*SD* = 0.25 dva) and those having larger eccentricities (*≥* 3 dva) by random Gaussian noise with a larger standard deviation (*SD* = 2 dva). **B.** The same as in Figure 4, although here, we simulated a true effect, that is, a radial increase in eccentricity of 2 dva in the Interest as compared to the Baseline condition. **C.** The same as in Figure 4, although here, equidistant binning was used. The equidistant bins ranged from an eccentricity of 0 dva to an eccentricity of 19.25 dva with a constant bin-width of 1.75 dva. Please note the different number of bins here relative to the other figure panels (11 vs 10, respectively).

Critically, both true effects and equidistant binning can substantially modify the appearance of the regression artifact. Along with circular condition cross-thresholding and eccentricity-dependent noise, this teaches us an important lesson: the regression artifact can take pretty much *any* form^8^.

For all presented simulation cases (null effect, null effect with cross-thresholding or eccentricity-dependent noise, and true effect), the regression artifact likewise manifested for another kind of binning analysis, namely, when binning the *x*_0_ and *y*_0_ values according to both eccentricity and polar angle (i.e., 2D segments) and computing shift vectors (Figure 2 as well as Figure 7 and Figure S2-S5, 1^st^ row). Here, the bin-wise means regressed towards and away from the overall means of the *x*_0_ and *y*_0_ distribution. The calculation of shift vectors is not uncommon in pRF studies (e.g., Klein et al., 2014; van Es et al., 2018; Vo et al., 2017).

**Figure 7.**
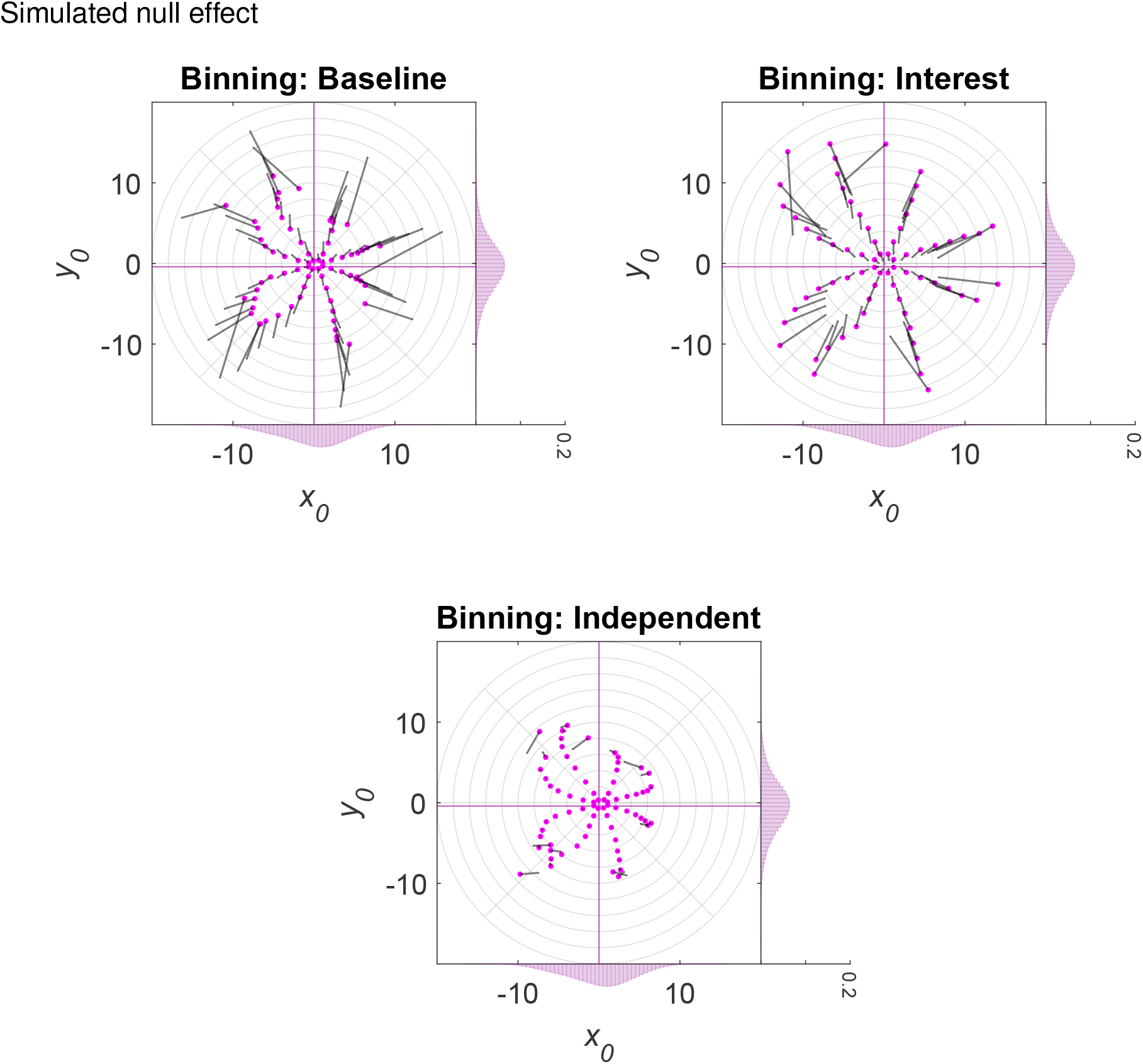
Simulated 2D post hoc binning analysis on *x*_0_ and *y*_0_ | Null effect. Bin-wise *x*_0_ and *y*_0_ means in the Interest and Baseline condition for a simulated null effect and different data binning scenarios. The *x*_0_ and *y*_0_ values in the Baseline and Interest condition were either binned according to eccentricity and polar angle values in the Baseline (1^st^ column, 1^st^ row), Interest (2^nd^ column, 1^st^ row), or an Independent condition (equivalent to repeat data of the Baseline condition; 2^nd^ row). The marginal histograms (bin width = 0.5 dva; *y*-axis: relative frequency) show the simulated *x*_0_ and *y*_0_ distributions for each condition, obtained by repeatedly disturbing the *x*_0_ and *y*_0_ values of an empirical visual field map with random Gaussian noise (*SD*= 2 dva). Magenta histograms correspond to the Interest condition and gray histograms to the Baseline condition. Note that the range of the marginal *y*-axis is the same for all histograms. The large magenta dots (arrow tip) correspond to the means in the Interest condition and the endpoint of the gray line (arrow knock) to the means in the Baseline condition. The gray line itself (arrow shaft) depicts the shift from the Baseline to the Interest condition. The magenta crosshair indicates the location of the overall *x*_0_ and *y*_0_ means for the Interest condition and the gray crosshair the location of the overall means for the Baseline condition. Note that if there is no systematic difference between the Baseline and Interest condition, the histograms and crosshairs coincide (as is the case here). The light gray polar grid demarks the bin segments. Polar angle bins ranged from 0° to 360° with a constant bin width of 45° and eccentricity bins from 0 to 20 dva with a constant bin width of 2 dva. The maximal eccentricity of the stimulated visual field area subtended 8.5 dva. Dva = Degrees of visual angle.

Notably, for empirical repeat data from the HCP (Benson et al., 2018, 2020), both kinds of binning analyses produced patterns consistent with the regression artifact (Figure S6-S13). This establishes its practical relevance. Moreover, some of us recently retracted an article on attention-induced differences in pRF position and size in V1-V3 (de Haas et al., 2014) because an in-house reanalysis suggested that circular post hoc binning along with circular condition cross-thresholding and heteroskedasticity yielded artifactual results in the form of egression from the mean (de Haas et al., 2020). In this case, the apparent significant effect was an increase in eccentricity and pRF size in the Interest vs Baseline condition (expressed as change scores) for eccentricity bins (based on the Baseline condition) in the middle of the tested range. Importantly, the inferential statistical analysis in this study (de Haas et al., 2014, 2020) was based on unbinned data, and thus the overall means. As such, the apparent significant effect was likely driven by or inflated due to circular cross-thresholding.

The example of de Haas et al. (2014, 2020) illustrates that data visualizations and associated inferential statistical analyses do not necessarily suffer from the same pitfalls. It is also possible that only one but not the other produces artifactual changes. This potential divergence adds another layer of complexity to the issues we discussed here.

Taken together, the heterogeneity in manifestation we exposed here makes it hard to spot the regression artifact by visual inspection alone and highlights its dependency on the type of analysis, additional circular selection practices as well as exact distributional properties of the data at hand (see Campbell and Kenny, 1999; Holmes, 2009; Schwarz and Reike, 2018, for similar points). Importantly, circular data binning is only but one pitfall resulting in artifactual changes. Other pitfalls, such as circular sorting of change scores and circular cross-thresholding are equally problematic.

### 3.2. Potential mitigation strategies

How can we omit double-dipping and control for regression towards the mean? We could, for instance, use an Independent condition for binning (such as repeat data or odd or even runs for the Baseline condition; Figure 4 and Figure 5-6, A.-C., 3^rd^ column as well as Figure 7 and Figure S2-S5, 2^nd^ row) or an anatomical criterion (Kriegeskorte et al., 2009), such as cortical distance or anatomical atlases (Benson et al., 2012, 2014). This way, noise components should be unbiased on average in both the Baseline and Interest condition.

Unbiased bin-wise noise components are of course less likely for sparsely populated bins (Figure 6, C., 3^rd^ column as well as Figure 7 and Figure S2-S5, 2^nd^ row), which can be captured by quantifying uncertainty. Critically, however, for scatter plots of change scores, bin-wise noise components are not unbiased for the Independent binning condition (Figure 5, A., 3^rd^ column). The reason for this is the same as before: non-independence of noise components. Thus, only the bin-wise change scores can be readily interpreted here. Moreover, given that cross-thresholding reshapes noise components, they might not be unbiased when binning on an Independent condition (Figure 5, B. and C., 3^rd^ column as well as Figure S2-S3, 2^nd^ row). The same can evidently also happen with an anatomical criterion if the Baseline and/or the Interest condition are subjected to cross-thresholding. Consequently, unless cross-thresholding can be omitted or demonstrated to be unbiased (see below for further considerations), an Independent condition might not be a safe option.

Of note, for the discussed cross-thresholding case where circular cross-thresholding was performed based on both the Interest and Baseline condition, binning on the Independent condition ensured that the bin-wise noise components for the Interest and Baseline condition are similarly biased (Figure 5, C., 3^rd^ column). As mentioned earlier, this is because cross-thresholding of this sort biases the noise components in the Baseline and Interest condition similarly (3.1. The many faces of regression towards the mean and other problems) and binning on an Independent condition introduces no further biases. Moreover, given that the noise components of both the Interest and Baseline condition were independent of those in the Independent condition, cross-thresholding did not bias the noise components in the Independent condition. As such, although the simple bin-wise means in the Baseline and Interest condition are biased, the difference between those amounts to around 0 (Figure 5, C., 3^rd^ column).

Apart from binning on an Independent condition, we could use analyses without binning that control for circularity and regression artifacts or effects could be evaluated against appropriate null distributions that take into account all statistical dependencies (e.g., Holmes, 2009; Kriegeskorte et al., 2009). For instance, errors-in-variables models (e.g., Deming regression) might be an option. Such models account for the noise in both the Baseline and Interest condition as well as for the fact that we often have no clear separation between independent and dependent variables in post hoc analyses of pRF data. However, as with any statistical approach, the underlying assumptions need to be checked carefully.

Just like circular data binning, circular sorting of change scores can be counteracted by plotting individual change scores against an Independent condition (Figure 5, A., 3^rd^ column). Similarly, one way to deal with circular cross-thresholding might be to cross-threshold all data according to an Independent condition/the Independent binning condition. However, condition-specific systematic errors, such as artifacts and outliers, might survive such independent data cleaning. As such, the usage of robust estimators might be advisable. Future research is necessary to evaluate this point more comprehensively.

A combination of the discussed approaches might prove most fruitful. Regardless of the specific mitigation strategy, we believe that in light of the many layers of complexity in our analysis pipelines, we need to make it common practice to perform sanity checks using (null) simulations and empirical repeat data. This is because such sanity checks provide a means for us researchers to ensure the validity of our analysis procedures.

### 3.3. The bigger picture

Circular post hoc binning analyses come in many flavors (e.g., centroids, shift vectors, eccentricity differences, *x*_0_ and *y*_0_ differences, and 1D or 2D bins) and cannot be assumed to be restricted to pRF position estimates. For instance, partial stimulation of pRFs likely results in heteroskedasticity and positively correlated errors for pRF size and position. This would, for instance, bias bin-wise pRF size vs pRF position or pRF size vs pRF size comparisons where binning is based on non-independent eccentricity values. Likewise, fitting errors due to partial stimulation should be more pronounced whenever pRF size is larger, leading to stronger artifactual effects (for simulations using different levels of noise see Holmes, 2009). The same is to be expected based on a higher variability in pRF position estimates for wider pRFs. These factors might potentially explain why changes in pRF position and/or size have been reported to be tendentially larger in higher-level areas where pRFs are wider (e.g., Barton and Brewer, 2015; de Haas et al., 2014, 2020; Klein et al., 2014; van Es et al., 2018).

Moreover, the distribution of errors likely depends on the toolbox that was used for fitting (Lerma-Usabiaga et al., 2020), making it hard to generalize across studies. And lastly, delineations of visual areas in post hoc binning analyses should ideally also be based upon independent criteria as this is where selection starts. Importantly, the intricacies we just discussed do not only apply to circular data binning, but also circular sorting of change scores and circular condition cross-thresholding.

The application of circular data binning, circular sorting of change scores, and/or circular cross-thresholding in the pRF literature might have led to spurious claims about changes in pRFs (see de Haas et al., 2014, 2020, for an example). Consequently, we encourage researchers who used such procedures to check for the severity of biases in their analyses by running adequate simulations and reanalyzing the original data wherever possible. Likewise, we urge them to take into account the issues discussed here when conducting future studies, reviewing manuscripts, and when teaching and mentoring.

### 3.4. Limitations

Our simulations were designed to encapsulate a given issue succinctly and cannot be interpreted as reflecting the exact properties of empirical pRF data. For this, we would need to have a good understanding of the underlying noise components. Similarly, the level of random Gaussian noise we adopted for most simulations (*SD* = 2 dva) might be more reminiscent of higher than lower visual areas (although this depends on many factors, such as mapping stimulus and magnetic field strength). For the present purposes, it appeared important to settle on a level allowing for clear exposition. Moreover, as alluded to further above (1. Introduction), unless there is a perfect correlation between two variables (and thus no random noise), double-dipping and/or regression towards or away from the mean likely pose issues to post hoc analyses involving a range of selection procedures, such as data binning, cleaning, and sorting.

To fully parallel our simulations, the analyses of the HCP data would have benefited from binning on an Independent condition, that is, a second set of repeat data. PRF estimates for such an Independent condition are currently not publicly available (Benson et al., 2018, 2020), leaving this sanity check for future research. Moreover, unlike our simulations, the condition cross-thresholding applied to the HCP data not only involved pRF position, but also goodness-of-fit (2.1. Post hoc binning using simulated data and 2.2. Post hoc binning using empirical repeat data). This is because such multivariate data cleaning is frequently applied in pRF studies. It is challenging to simulate these more complex scenarios and thus best addressed in a separate article.

Some post hoc binning analyses in the pRF literature are conducted in a hemifield-specific fashion, whereas others mirror observations across hemifields or quadrants. Our analyses do not capture these specificities. However, there is no reason to believe that they would alleviate the expression of the regression artifact. The primary component that might change when applying such procedures is the location of the overall mean and the shape of the data distribution and thus how exactly the artifact manifests (for preliminary analyses, see Stoll et al., 2022). Of course, if data points are not mirrored based on an Independent condition but, for instance, the Baseline condition, data mirroring in combination with post hoc binning and/or circular cross-thresholding might favor noise components in multiple ways. Importantly, circular data mirroring is also problematic for analyses that do not involve any circular data binning and/or circular cross-thresholding, as are other procedures, such as circular data weighting (Kriegeskorte et al., 2009).

## 4. Conclusions

Without doubt, circularity and regression towards the mean are thorny and omnipresent problems that can manifest subtly and diversely (e.g., Ball et al., 2020; Barnett et al., 2005; Campbell and Kenny, 1999; Eriksson and Häaggsträom, 2014; Gignac and Zajenkowski, 2020; Holmes, 2009; Kilner, 2013; Kriegeskorte et al., 2009; Preacher et al., 2005; Shanks, 2017; Stigler, 1997; Vul et al., 2009). As such, we need to ensure that the validation of analysis procedures becomes part and parcel of the scientific process.

## Data and code availability

Preprocessed data, custom code, and figures are available at https://doi.org/10.17605/OSF.IO/WJADP.

## Acknowledgements

This research was supported by European Research Council Starting Grants to DSS (WMOSPOTWU, 310829) and BdH (INDIVISUAL, 852885). BdH was further supported by the Deutsche Forschungsgemeinschaft (222641018–SFB/TRR 135 TP C9). We thank three peer reviewers for providing constructive feedback.

## Declaration of competing interest

The authors declare no conflict of interest.

## Supplementary methods

### 1. Retinotopic mapping experiment

#### 1.1. Participants

All participants (*N* = 5, of which 2 were authors; 3 females; age range: 29-36 years) had corrected-to-normal visual acuity (obtained through corrective contact lenses) and gave written informed consent. As mentioned in the main text (2.1. Post hoc binning using simulated data), only the dataset of a single participant was used for simulation purposes. Experimental procedures were approved by the University College London Ethics Committee.

#### 1.2. Apparatus

Functional and anatomical images were acquired at a field strength of 1.5 T on a Siemens Avanto magnetic resonance imaging (MRI) scanner. All stimuli were projected onto a screen (resolution: 1920 *×* 1080 pixels; refresh rate: 60 Hz; background color: gray) at the back of the MRI scanner. Participants viewed the experiment through a head-mounted mirror. The viewing distance was approximately 67 cm. To ensure unobstructed view, we used a custom-made 32-channel head coil, where the front visor was demounted, leaving 30 effective channels. Eye movements of participant’s left eye were recorded via an EyeLink 1000 MRI compatible eye tracker.

#### 1.3. Stimuli and procedure

The mapping stimulus comprised a gray square field with a dynamic horizontal bar aperture (length of major axis: 17.15 dva; length of minor axis: 1.27 dva). The bar aperture was presented within the boundaries of a circular mapping area (diameter: 17.15 dva). It moved discretely and consecutively across the mapping area along cardinal (0/180° and 90/270°) and oblique axes (45/225° and 135/315°) and was superimposed onto a random dot kinematogram (RDK). The RDK comprised moving black dots (diameter: 0.13 dva) positioned within a square field (size: 17.03 *×* 17.03 dva). If a dot left the square field, it was moved back by 1 field width/height. The dots had a density of 6.89 dots/dva^2^, a lifetime of 36 frames, were repositioned randomly once they had died, and oscillated coherently along the major axis of the bar aperture according to a sine wave (*A* = 1.29 dva, *f* = 1 Hz, *ω* = 6.28 rad/s, *ϕ* = 0 rad). The mapping stimulus and RDK were centered at the screen’s midpoint.

A semi-transparent (*α* = 50%) array of 5 vertical ovals was superimposed onto the mapping stimulus. One of the ovals was centered at the screen’s mid-point (length of major axis: 0.43 dva; length of minor axis: 0.28 dva) and the remaining ovals at an eccentricity of 4.29 dva (length of major axis: 0.86 dva; length of minor axis: 0.57 dva) and different polar angles (45°, 135°, 225°, and 315°). The ovals were presented as a rapid serial visual presentation (RSVP) task, where each trial started with 200 ms of oval presentation, followed by an interval of 600 ms without any ovals. Each oval’s orientation (45° left- or rightwards from vertical) and color (red, yellow, cyan, orange, brown, white, black, green, and blue) changed randomly in each trial with the exception that ovals of the same color were never presented simultaneously. Participants had to press a button whenever a rightwards oriented oval was presented in blue or green color. A black polar grid (line width: 0.02 dva) at low opacity (*α* = 20%) with 12 radial lines (polar angles: 0 to 330° with a step size of 30°) and 18 circles (diameters: 0.95 to 51.42 dva with a step size of 2.97 dva) was superimposed onto the screen. The radial lines ran from the midpoint of the screen to the outermost circle.

The experiment comprised 4 attention conditions, in which participants were required to perform the RSVP task on different oval streams whilst ignoring other streams and the bar aperture. The condition used for simulation purposes was the *Center* condition, where participants performed the task on the central oval stream. This condition therefore resembled a standard pRF mapping experiment where participants typically perform a task at fixation (e.g., Alvarez et al., 2015; Amano et al., 2009; Benson et al., 2018). Participants performed 2 sessions each with 4 runs per condition on consecutive days. The order of conditions was pseudorandomized. Participants’ eye position and pupil size were recorded at 60 Hz (down-sampled) throughout each run. One day prior to the first session, participants underwent 1 mock run per condition inside the scanner to familiarize themselves with the task. Here, only behavioral data (and no functional or anatomical images) were collected.

Within each run, the bar aperture moved along each axis twice, so that the starting point covered all chosen polar angles. Specifically, the sequence of starting points in each run was: 90°, 225°, 180°, 315°, 270°, 45°, 0°, and 135°. One bar sweep lasted 28 s (1 step/s). Consecutive bar apertures overlapped by 50%. After 4 bar sweeps, a blank interval of 28 s (without the bar apertures and RDK) was presented, during which participants had to refrain from doing the RSVP task. A brief tone cued the beginning and end of this interval. The position and lifetime of each dot in the RDK at the start of every 28-s-interval was randomized. Experimental procedures were implemented in Matlab 2014a (8.3; https://uk.mathworks.com/) using Psychtoolbox-3 (3.0.11; Brainard, 1997; Kleiner et al., 2007; Pelli, 1997).

#### 1.4. MRI acquisition

We collected anatomical images using a T1-weighted magnetization-prepared rapid acquisition with gradient echo sequence (repetition time, TR = 2.73 s; echo time, TE = 3.57 ms; voxel size = 1 mm isotropic; flip angle = 7°; field of view, FoV = 256 mm *×* 224 mm; matrix size = 256 *×* 224; 176 sagittal slices) and functional images using a T2*-weighted multiband 2D echo-planar imaging sequence (Breuer et al., 2005, TR = 1 s, TE = 55 ms, voxel size = 2.3 mm isotropic, flip angle = 75°, FoV = 224 mm × 224 mm, no gap, matrix size: 96 × 96, acceleration = 4, 36 transverse slices). The slab for the functional images was aligned to be roughly parallel to the calcarine sulcus so that the posterior third of the cortex was well covered.

#### 1.5. Preprocessing

The initial 10 volumes of each run were discarded to allow for magnetization to reach equilibrium. Using SPM8 (6313; https://www.fil.ion.ucl.ac.uk/spm/software/spm8/), functional images were then bias-corrected, realigned, unwarped, coregistered to the anatomical image, and finally projected onto an anatomical surface model constructed in FreeSurfer (5.3.0; Dale et al., 1999; Fischl et al., 1999). We generated vertex-wise fMRI time series per run by determining the functional voxel at half the distance between corresponding vertices in the pial surface and gray-white matter mesh. We then applied linear detrending to the time series of each run and *z*-standardized them. Surface projection, detrending, and *z*-standardization were performed in Matlab 2016b (9.1; https://uk.mathworks.com/) using SamSrf7 (7.05; https://github.com/samsrf/samsrf/tree/3c7a0e25090e9097d5e2fd95 696c00774acd26d6).

#### 1.6. pRF estimation and delineations

The vertex-wise preprocessed time series of the Center condition were averaged across the 2 sessions. We then fit a 2D isotropic Gaussian pRF model with 5 free parameters (*x*_0_, *y*_0_, *σ*, *β*_0_, and *β*_1_) to the vertex-wise average time series. To this end, we first predicted pRF responses by calculating the overlap between the pRF model and an indicator function of the bar aperture for each volume using a 100 *×* 100 pixel matrix. Specifically, we used a 3D search space of possible values for *σ* (8.5 dva *×* 2^-5.6:0.2:1^)^9^, *x*_0_, and *y*_0_, and generated pRF responses for each combination of these values. Values for *x*_0_ and *y*_0_ were first sampled from the polar coordinate system (polar angles: 0:10:350°; eccentricities: 8.5 dva *×* 2^-5:0.2:0.6^) and then transformed to Cartesian coordinates. The pRF response per volume was expressed as mean percent overlap with the pRF model.

To obtain a predicted fMRI time series, we then convolved these pRF responses with a canonical hemodynamic response function (HRF) obtained based on data from a previous study (de Haas et al., 2014, 2020). Next, we calculated the Pearson correlation between the predicted and the observed fMRI time series and retained the combination of parameter values showing the largest *R*^2^ with all *R*^2^s *≥* .01. These initial parameter estimates were then used as seeds for an optimization procedure aimed at further maximizing the Pearson correlation between the observed and predicted fMRI time series using a Nelder-Mead algorithm (Lagarias et al., 1998; Nelder and Mead, 1965). Lastly, we estimated *β*_0_ and *β*_1_ by performing linear regression between the observed and predicted time series. The final parameter maps were smoothed with a Gaussian kernel (FWHM = 3 mm) in spherical surface space. Vertices with a very poor *R*^2^ (*<* .01) or artifacts (*σ ≤* 0, *β*_1_ *≤* 0 or *β*_1_ *>* 3) were removed prior to smoothing. V1 hemifield maps were manually delineated based on smooth polar angle maps using polar angle reversals (Engel et al., 1997; Sereno et al., 1995; Wandell et al., 2007). These delineations were used as a mask to extract V1 vertices. Fitting, smoothing, and manual delineations were performed in Matlab 2016b (9.1; https://uk.mathworks.com/) using SamSrf7 (7.05; https://github.com/samsrf/samsrf/tree/3c7a0e25090e9097d5e2fd95696c00774acd26d6). The canonical HRF we adopted is implemented in SamSrf7.

## Supplementary figures

**Figure S1.**
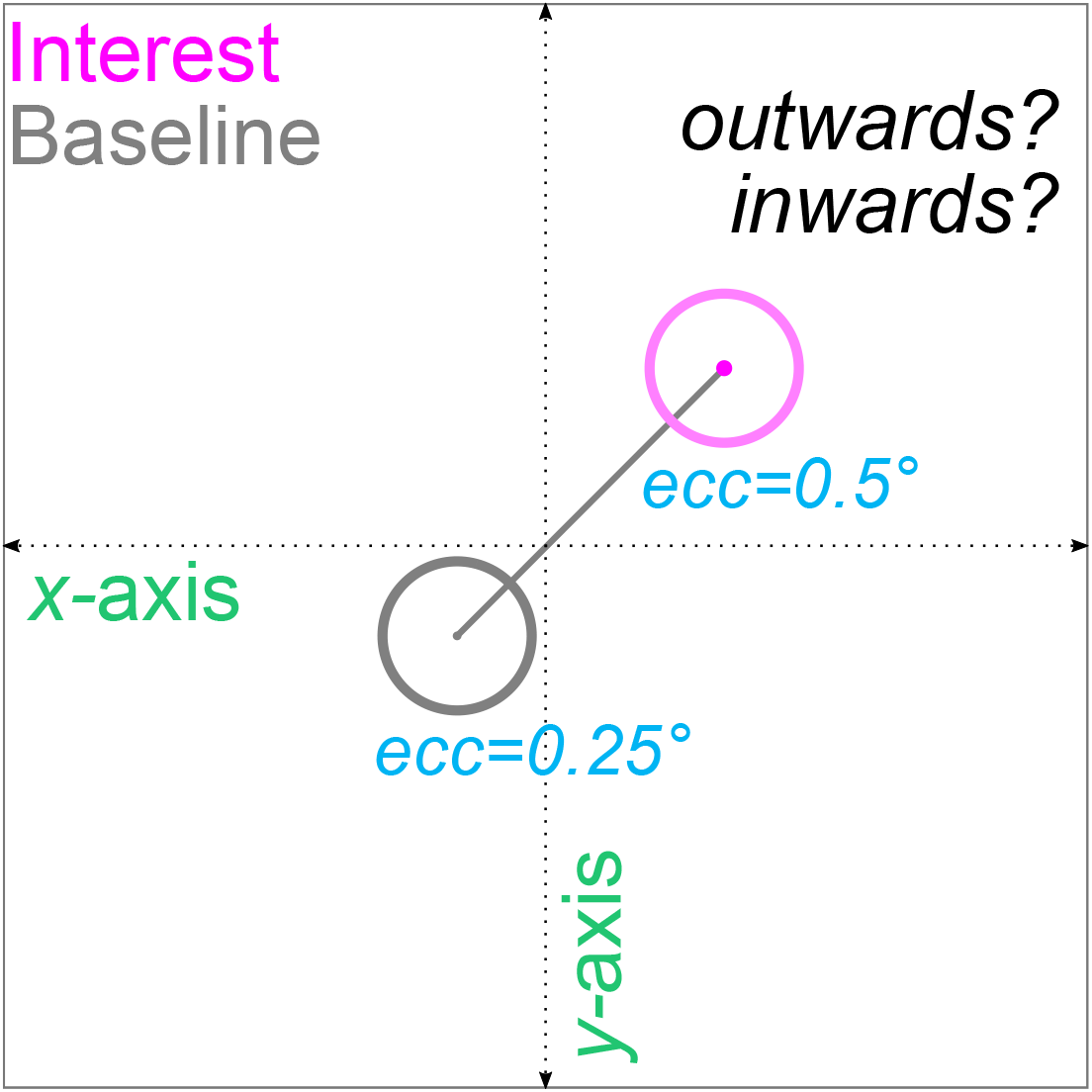
Interpretation of changes in eccentricity. The same as Figure 2, although here, the pRF shifts from one visual field quadrant to another in the Interest compared to the Baseline condition. This can happen due to noise or when visual field maps partially cover the ipsilateral hemifield. In such cases, an increase or decrease in eccentricity does not necessarily correspond to an outwards or inwards shift in the traditional sense. For instance, imagine that a pRF sits at *x*0 = -0.18 dva and *y*0 = -0.18 dva in the Baseline condition (ecc = 0.25 dva) but at *x*0 = 0.36 dva and *y*0 = 0.36 dva in the Interest condition (ecc = 0.51 dva). This would result in an increase in eccentricity, which might be interpreted as an outwards shift, although the pRF shifts effectively radially inwards until it reaches the origin and then outwards. We can likewise imagine that the pRF shifts horizontally to *x*0 = 0.36 dva and *y*0 = -0.36 dva in the Interest condition. Importantly, removing such shifts would bias noise components (see condition cross-thresholding in the main text and Figure 5, B. and C. as well as Figure S2-S3) and therefore, does not seem a valid option. Dva = Degrees of visual angle.

**Figure S2.**
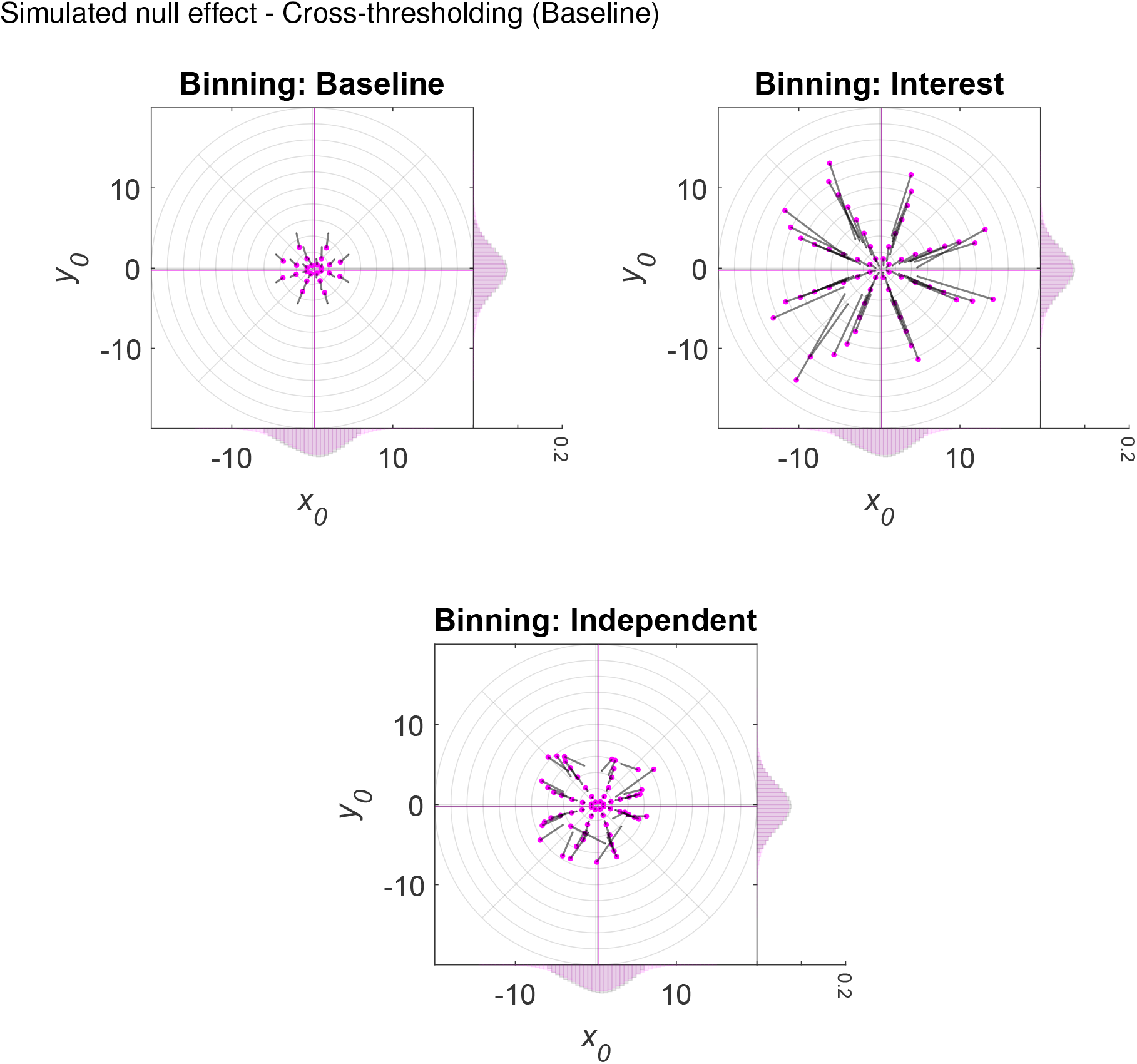
Simulated 2D post hoc binning analysis on *x*_0_ and *y*_0_ | Null effect — Cross-thresholding (Baseline). The same as in Figure 7, although here, condition cross-thresholding was applied, i.e., simulated observations falling outside a certain eccentricity range (*≥* 0 and *≤* 6 dva) in the Baseline condition were removed from all conditions. (Condition) cross-thresholding = The pair-wise or list-wise deletion of observations across conditions.

**Figure S3.**
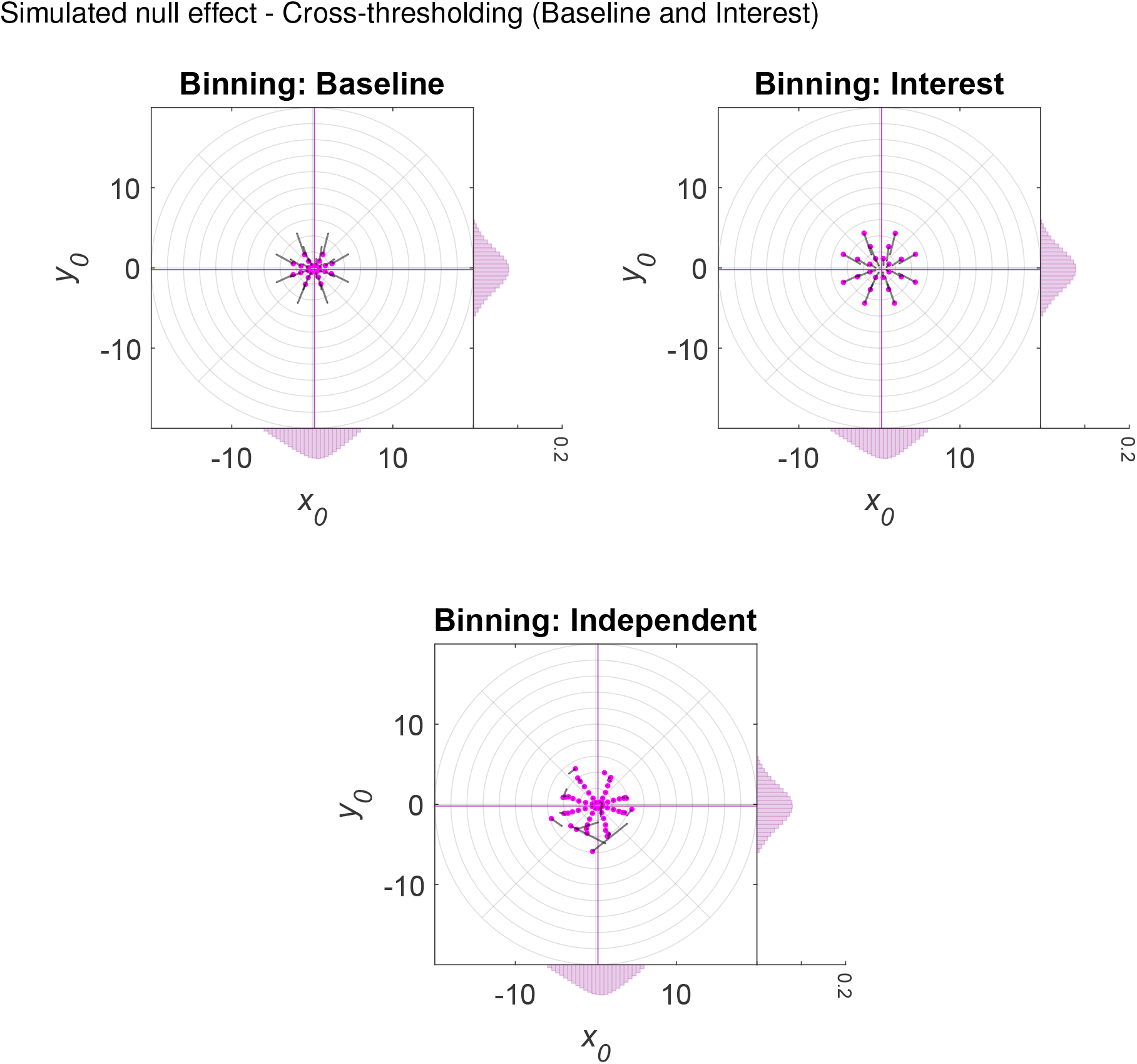
Simulated 2D post hoc binning analysis on *x*_0_ and *y*_0_ | Null effect — Cross-thresholding (Baseline and Interest). The same as in Figure S2, although here, condition cross-thresholding was based on both the Baseline and Interest condition.

**Figure S4.**
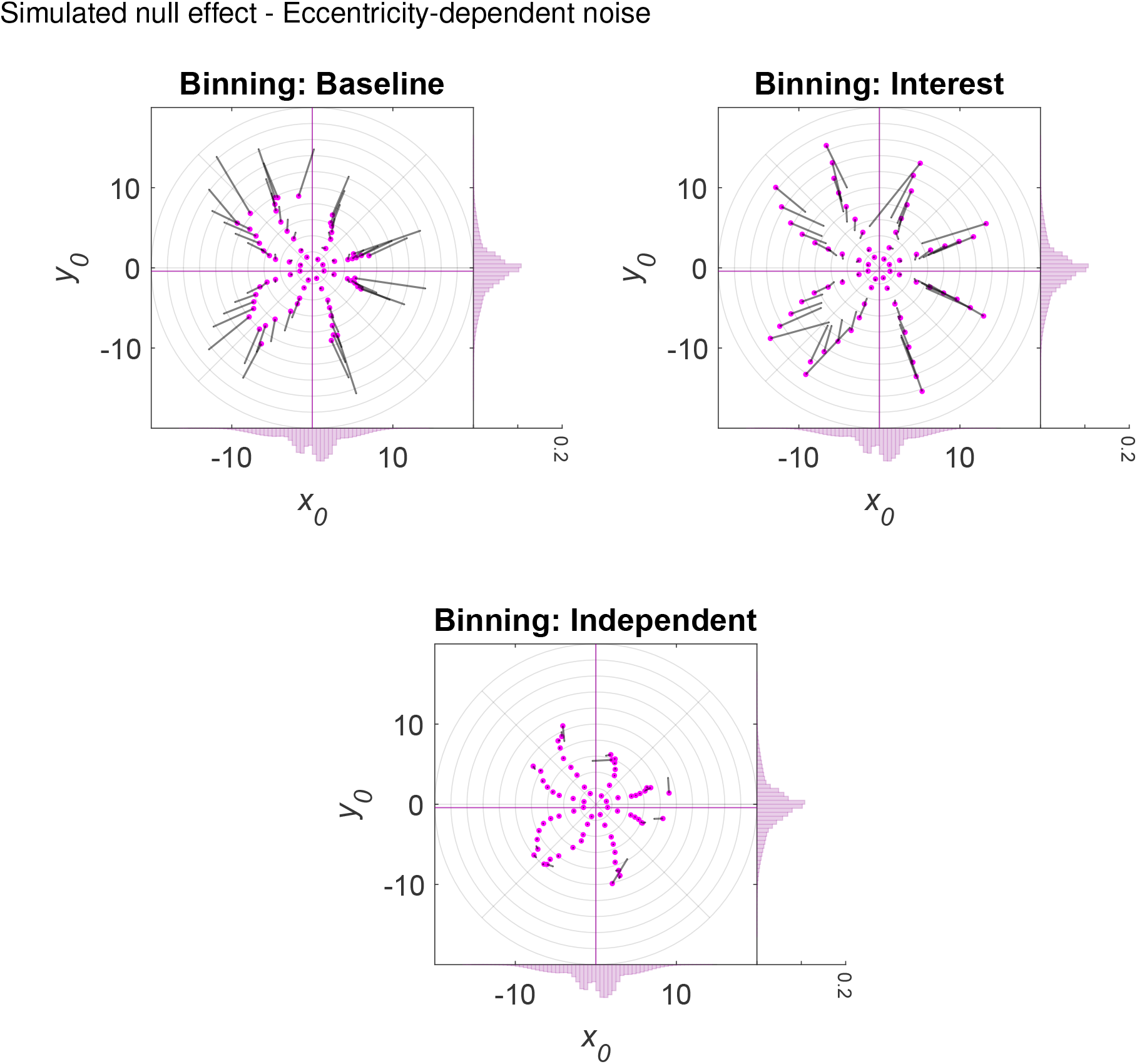
Simulated 2D post hoc binning analysis on *x*0 **and** *y*0 | Null effect — Eccentricity-dependent noise. The same as in Figure 7, although here, original observations having smaller eccentricities (*≥* 0 and *<* 3 dva) were disturbed by random Gaussian noise with a smaller standard deviation (*SD* = 0.25 dva) and those having larger eccentricities (*≥* 3 dva) by random Gaussian noise with a larger standard deviation (*SD* = 2 dva).

**Figure S5.**
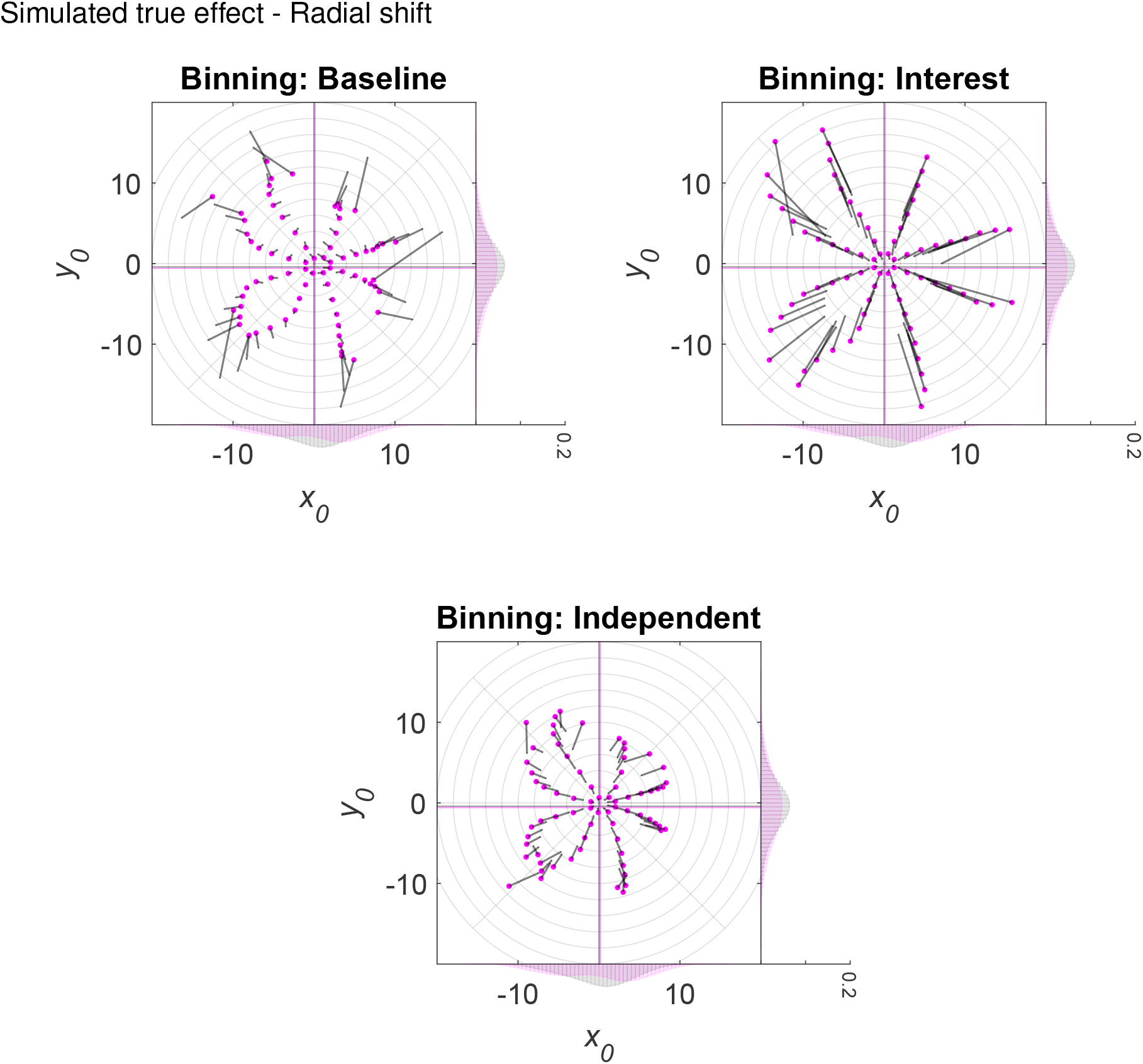
Simulated 2D post hoc binning analysis on *x*_0_ and *y*_0_ | True effect — Radial shift. The same as in Figure 7, although here, we simulated a true effect, that is, a radial increase in eccentricity of 2 dva in the Interest as compared to the Baseline condition. Note that the eccentricity bins ranged from 0 to 22 dva here (instead of 0 to 20 dva).

**Figure S6.**
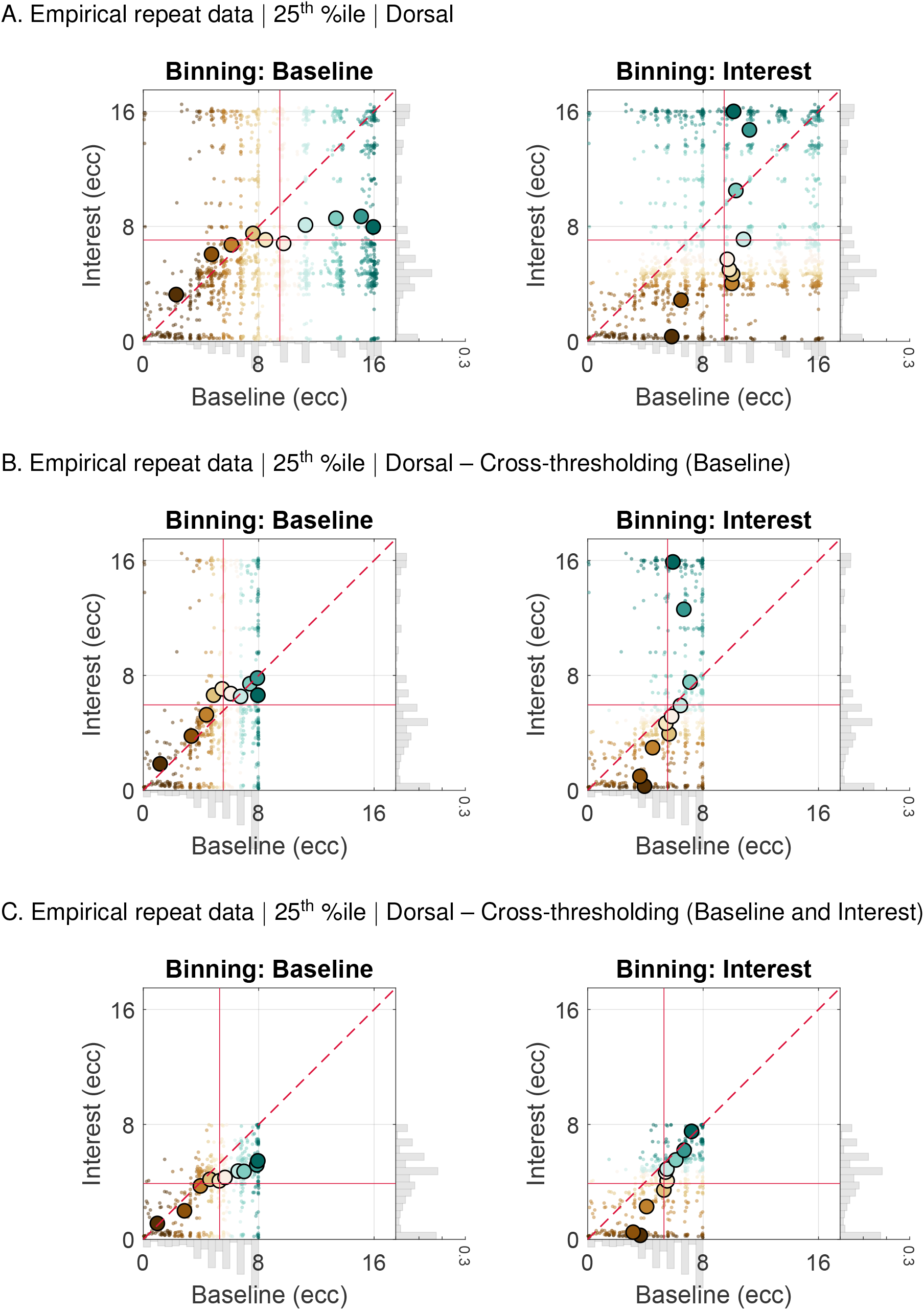
Empirical 1D post hoc binning analysis on eccentricity | Repeat data | 25^th^ %ile participant | Dorsal — Without and with cross-thresholding. Bin-wise eccentricity values and means in the Interest and Baseline condition for repeat data from the HCP belonging to the 25^th^ %ile participant of the median *R*^2^ distribution and different data binning scenarios. **A.** Data from the dorsal complex (V3A/B and IPS0–5) without condition cross-thresholding. **B.** Same as A., but with condition cross-thresholding. To this end, eccentricity values falling outside a certain eccentricity range (*≥* 0 and *≤* 8 dva) and below a certain *R*^2^ cut-off (*≤* 2.2%) in the Baseline condition were removed from both conditions. **C.** Same as B., although here, condition cross-thresholding was based on both the Baseline and Interest condition. The eccentricity values in the Baseline and Interest condition were either binned according to eccentricity values in the Baseline (1^st^ column in A.-C.) or Interest (2^nd^ column in A.-C.) condition. The gray marginal histograms (bin width = 0.5 dva; *y*-axis: relative frequency) show the eccentricity distributions for each condition. Note that the range of the marginal *y*-axis is the same for all histograms. The red crosshair indicates the location of the overall mean for the Interest and Baseline condition. The red dashed line corresponds to the identity line. Dark brown colors correspond to lower and dark blue-green colors to higher decile bins. The smaller colorful dots represent individual data points and the larger colorful dots with the black outline bin-wise means. The maximal eccentricity of the stimulated visual field area subtended 8 dva. HCP = Human Connectome Project. Dva = Degrees of visual angle. Ecc = Eccentricity. %ile = Percentile. (Condition) cross-thresholding = The pair-wise or list-wise deletion of observations across conditions.

**Figure S7.**
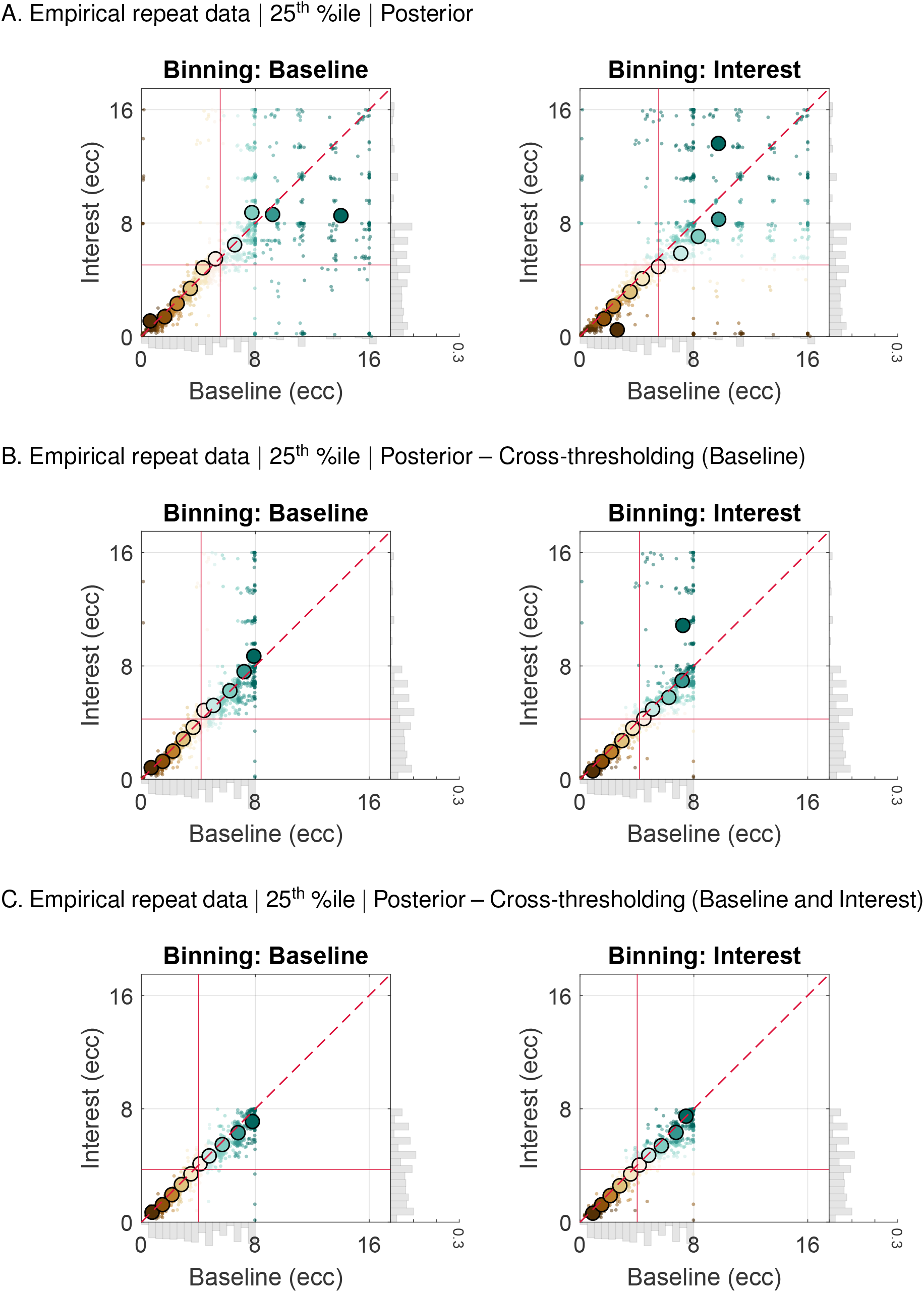
Empirical 1D post hoc binning analysis on eccentricity | Repeat data | 25^th^ %ile participant | Posterior — Without and with cross-thresholding. The same as in Figure S6, although here, we used data from the posterior complex (V1-V3).

**Figure S8.**
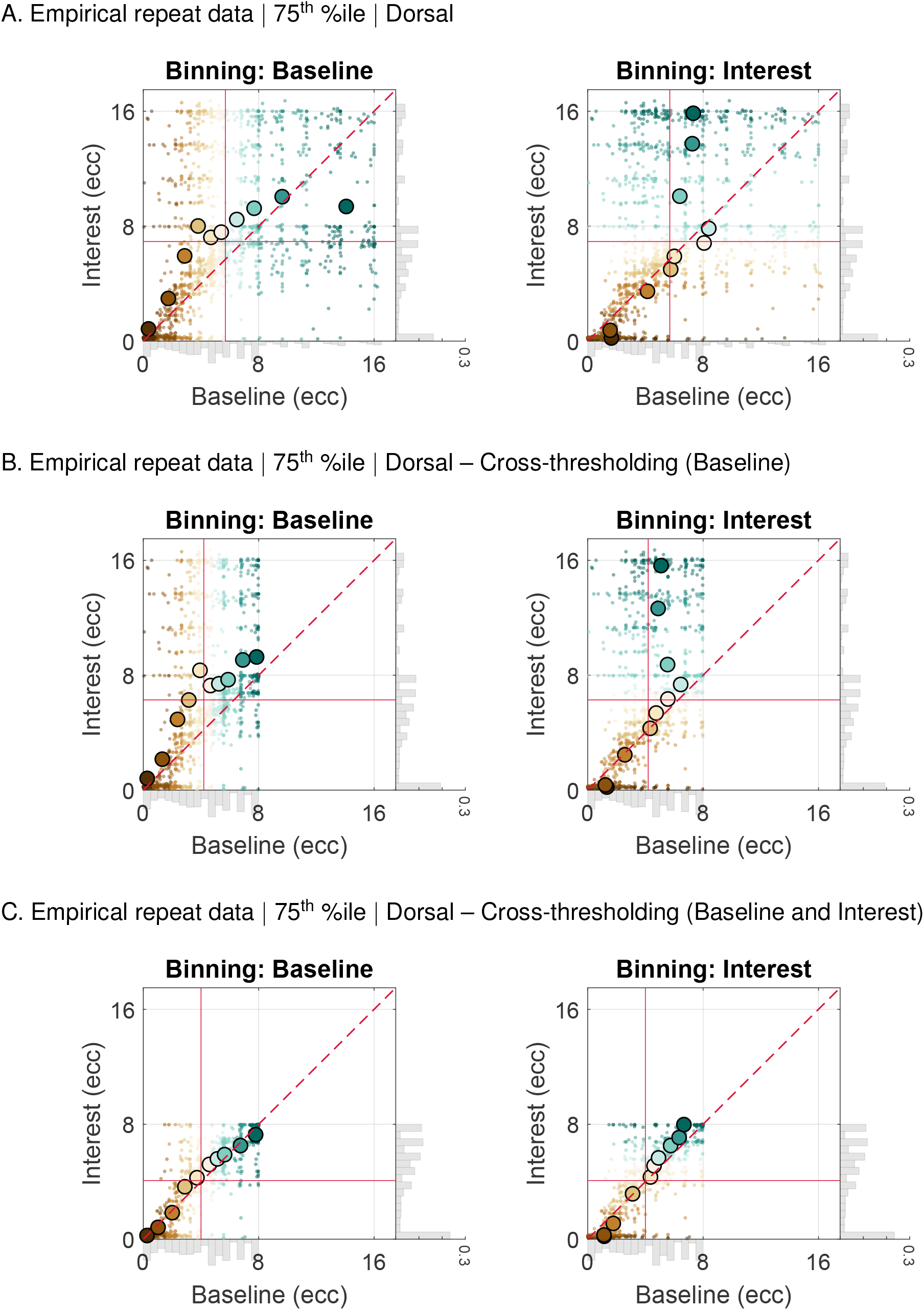
Empirical 1D post hoc binning analysis on eccentricity | Repeat data | 75^th^ %ile participant | Dorsal — Without and with cross-thresholding. The same as in Figure S6, although here, we used the 75^th^ %ile participant of the median *R*^2^ distribution.

**Figure S9.**
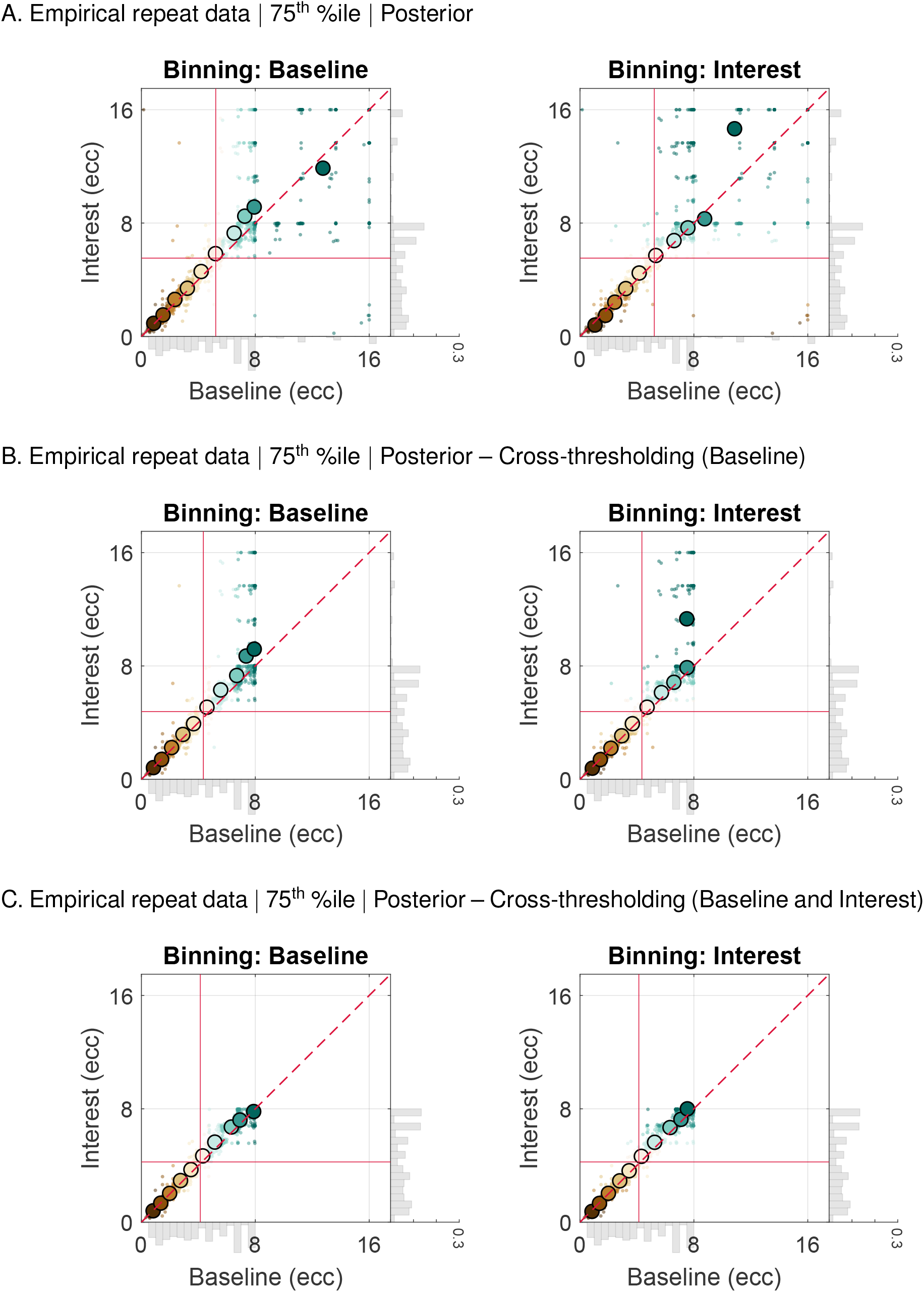
Empirical 1D post hoc binning analysis on eccentricity | Repeat data | 75^th^ %ile participant | Posterior — Without and with cross-thresholding. The same as in Figure S7, although here, we used the 75^th^ %ile participant of the median *R*^2^ distribution.

**Figure S10.**
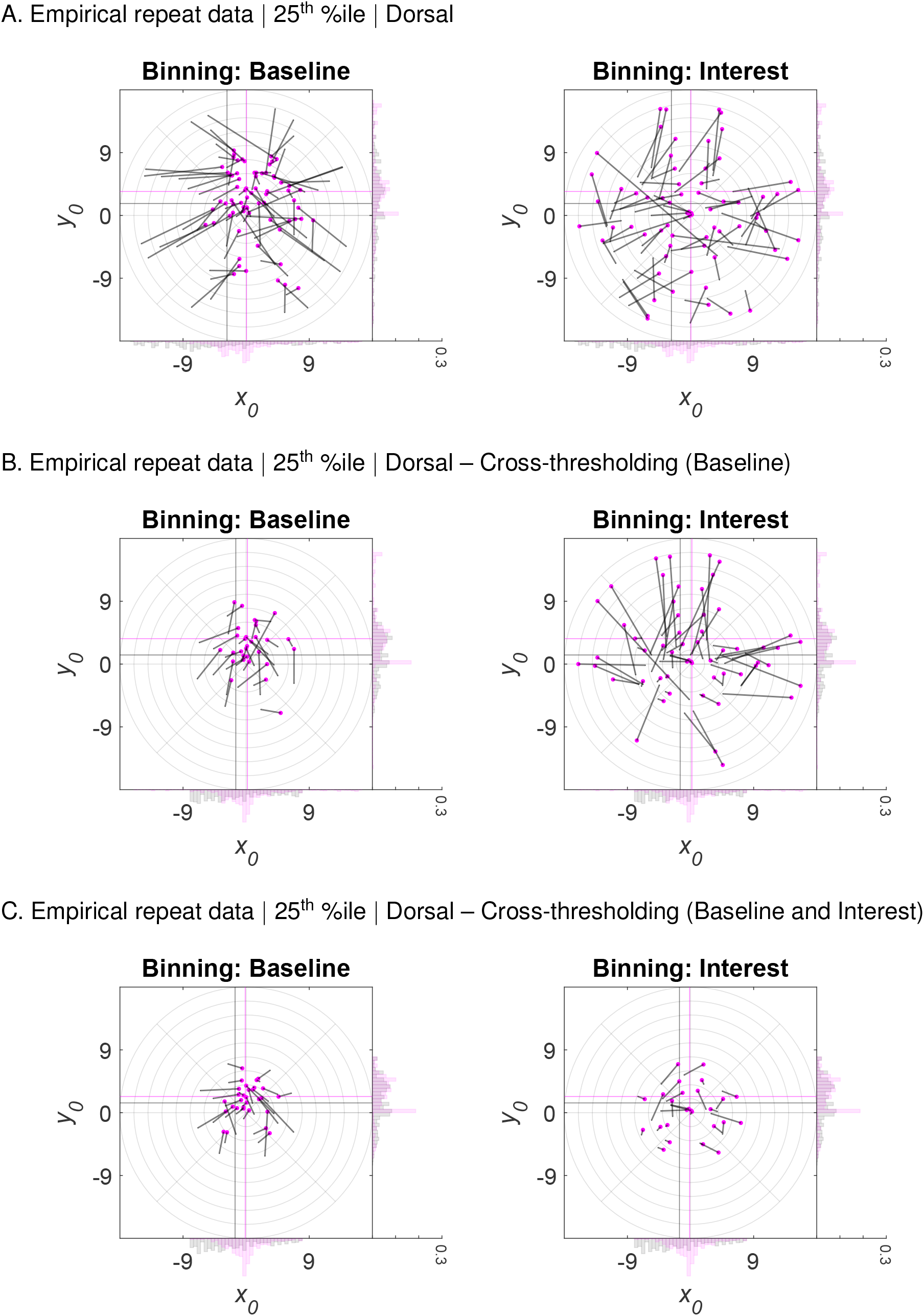
Empirical 2D post hoc binning analysis on *x*_0_ and *y*_0_ | Repeat data | 25^th^ %ile participant | Dorsal — Without and with cross-thresholding. Bin-wise *x*_0_ and *y*_0_ means in the Interest and Baseline condition for repeat data from the HCP belonging to the 25^th^ percentile participant of the median *R*^2^ distribution and different data binning scenarios. **A.** Data from the dorsal complex (V3A/B and IPS0–5) without condition cross-thresholding. **B.** Same as A., but with condition cross-thresholding. To this end, eccentricity values falling outside a certain eccentricity range (*≥* 0 and *≤* 8 dva) and below a certain *R*^2^ cut-off (*≤* 2.2%) in the Baseline condition were removed from both conditions. **C.** Same as B., although here, condition cross-thresholding was based on both the Baseline and Interest condition. The *x*_0_ and *y*_0_ values in the Baseline and Interest condition were either binned according to eccentricity and polar angle values in the Baseline (1^st^ column in A.-C.) or Interest (2^nd^ column in A.-C.) condition. The marginal histograms (bin width = 0.5 dva; *y*-axis: relative frequency) show the *x*_0_ and *y*_0_ distributions for each condition. Magenta histograms correspond to the Interest condition and gray histograms to the Baseline condition. Note that the range of the marginal *y*-axis is the same for all histograms. The large magenta dots (arrow tip) correspond to the means in the Interest condition and the endpoint of the gray line (arrow knock) to the mean in the Baseline condition. The gray line itself (arrow shaft) depicts the shift from the Baseline to the Interest condition. The magenta crosshair indicates the location of the overall *x*_0_ and *y*_0_ means for the Interest condition and the gray crosshair the location of the overall means for the Baseline condition. Note that for subtle differences between the Baseline and Interest condition, the histograms and crosshairs almost coincide (see Figure S11 and Figure S13). The light gray polar grid demarks the bin segments. Polar angle bins ranged from 0° to 360° with a constant bin width of 45° and eccentricity bins from 0 to 18 dva with a constant bin width of 2 dva. The maximal eccentricity of the stimulated visual field area subtended 8 dva. HCP = Human Connectome Project. Dva = Degrees of visual angle. %ile = Percentile. (Condition) cross-thresholding = The pair-wise or list-wise deletion of observations across conditions.

**Figure S11.**
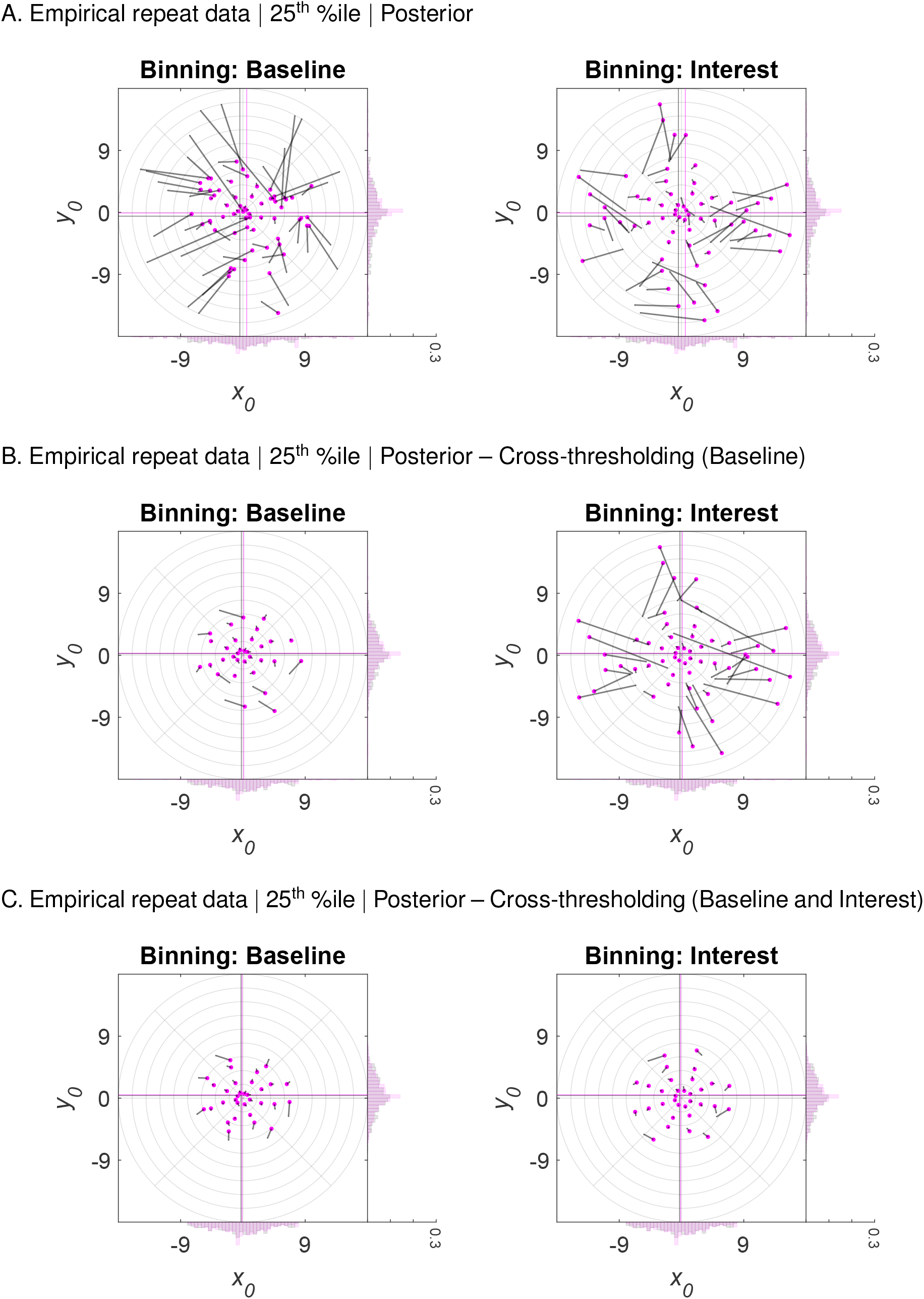
Empirical 2D post hoc binning analysis on *x*_0_ and *y*_0_ | Repeat data | 25^th^ %ile participant | Posterior — Without and with cross-thresholding. The same as in Figure S10, although here, we used data from the posterior complex (V1-V3).

**Figure S12.**
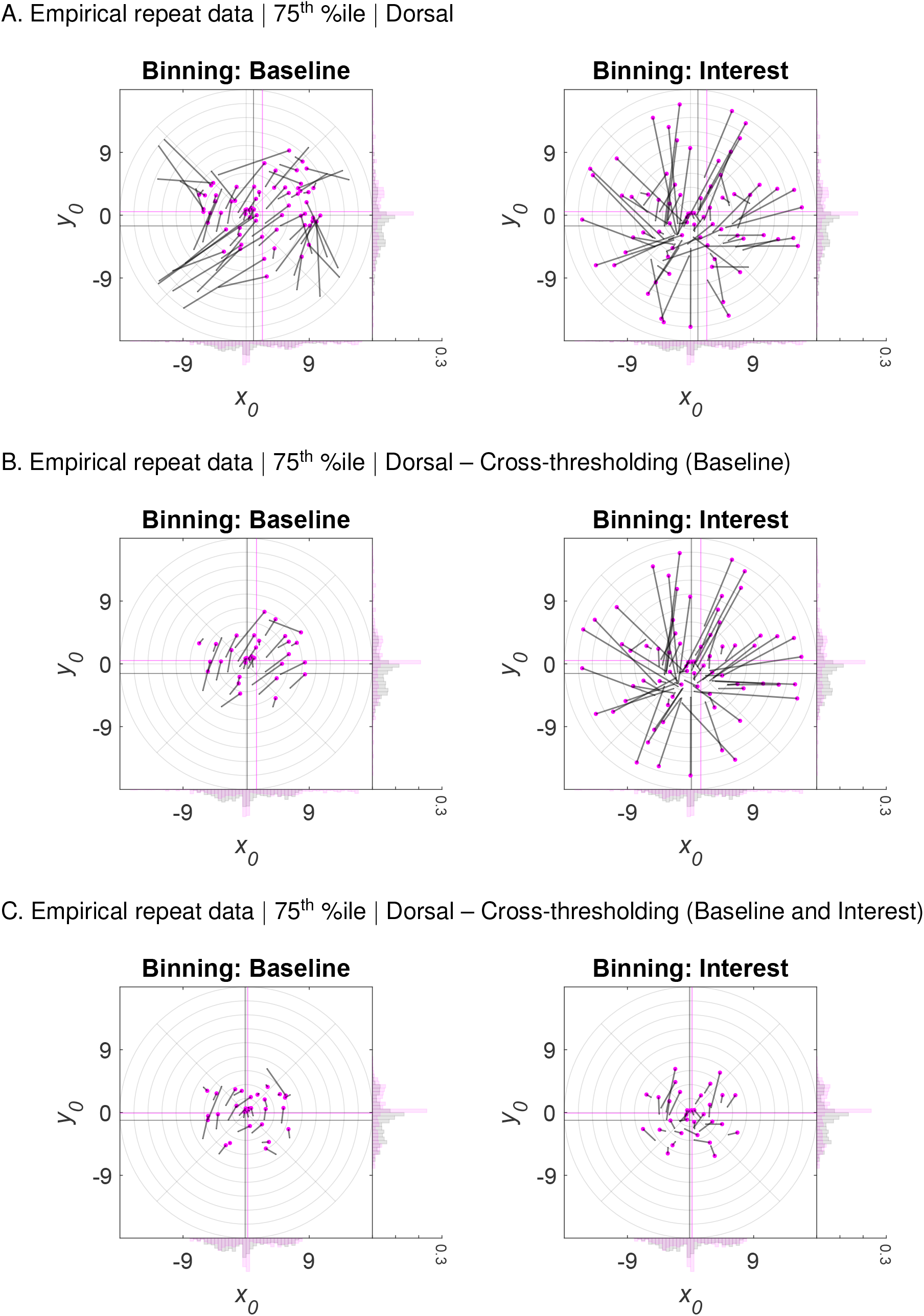
Empirical 2D post hoc binning analysis on *x*_0_ and *y*_0_ | Repeat data | 75^th^ %ile participant | Dorsal — Without and with cross-thresholding. The same as in Figure S10, although here, we used the 75^th^ %ile participant of the median *R*^2^ distribution.

**Figure S13.**
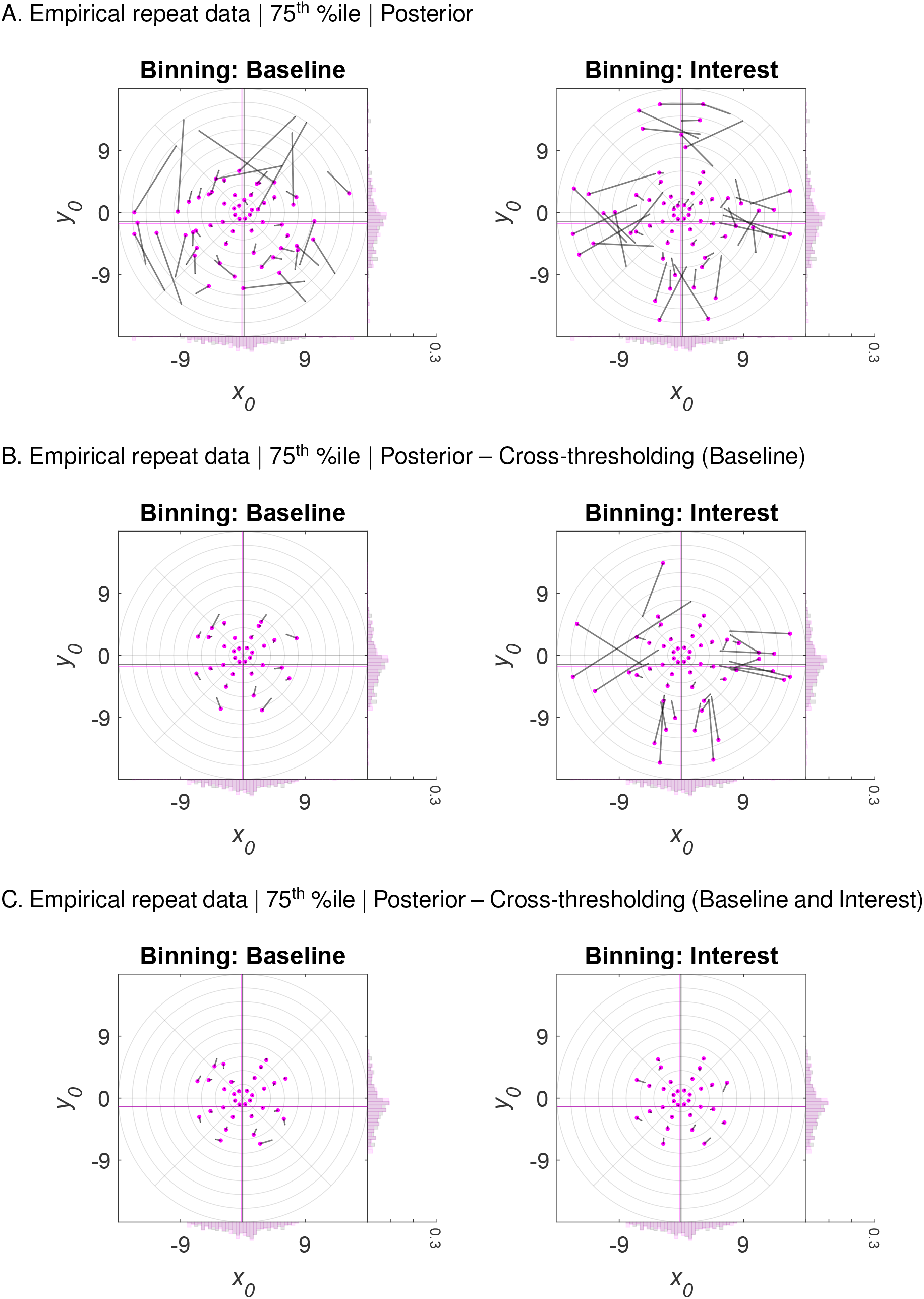
Empirical 2D post hoc binning analysis on *x*_0_ and *y*_0_ | Repeat data | 75^th^ %ile participant | Posterior — Without and with cross-thresholding. The same as in Figure S11, although here, we used the 75^th^ %ile participant of the median *R*^2^ distribution.

## Footnotes

1 Note that random noise is only one factor weakening the correlation between two variables (for more details, see Shanks, 2017).

2 To be precise, regression towards the mean refers to standard scores (*z*-scores; Campbell and Kenny, 1999; Kenny, 2005)

3 Note that when evaluating data distributions with unequal means, variances, or non-linearity, *z*-standardization might be necessary to detect regression towards or away from the mean (Campbell and Kenny, 1999; Shanks, 2017). In particular, *z*-standardization makes data distributions directly comparable. As such, bin-wise means should regress to wherever they intersect the identity line. Here, we always display data in native space, as this is typically done in the pRF literature. However, we use crosshairs to indicate the location of the mean and thus provide a visual guideline.

4 Note that apart from the visualizations provided here, it might be beneficial to additionally look at Galton squeeze diagrams to detect regression towards or away from the mean (see Figure 1; Campbell and Kenny, 1999; Galton, 1886; Shanks, 2017)

5 Note that for skewed distributions (such as the gamma-like distribution here), the regression effect might be actually towards the mode and away from the mean of the overall distribution (Schwarz and Reike, 2018). If the location of the overall mode and mean are sufficiently close, our visualizations would be unable to distinguish these two cases.

6 For reasons of clarity and simplicity, we use the term ‘circular condition’ or ‘non-circular condition’ exclusively when referring to circular data binning. However, other circular selection procedures, such as circular data sorting or cleaning, might of course render a condition circular above and beyond circular data binning.

7 Note that the regression was presumably towards the nearest modes of the simulated bimodal distribution (see marginal histograms in Figure 6, A., 1^st^ and 2 ^nd^ columns; Schwarz and Reike, 2018).

8 Note that floor/ceiling effects (due to physiological and methodological constraints on the minimum and maximum observable value) and/or the calculation of absolute (raw) vs proportional (%) differences are further factors influencing the appearance of the regression artifact (de Haas et al., 2014, 2020; Holmes, 2009).

9 Note that j:i:k stands for a regularly-spaced vector where i reflects the increment between j and k.

## Notes

### Competing Interest Statement

The authors have declared no competing interest.

### Summary of Updates

Figure 1 revised; several sections rewritten; several sections removed.

